# AlphaFold2-Enabled Atomistic Modeling of Epistatic Binding Mechanisms for the SARS-CoV-2 Spike Omicron XBB.1.5, EG.5 and FLip Variants: Convergent Evolution Hotspots Cooperate to Control Stability and Conformational Adaptability in Balancing ACE2 Binding and Antibody Resistance

**DOI:** 10.1101/2023.12.11.571185

**Authors:** Nishank Raisinghani, Mohammed Alshahrani, Grace Gupta, Sian Xiao, Peng Tao, Gennady Verkhivker

## Abstract

In this study, we combined AI-based atomistic structural modeling and microsecond molecular simulations of the SARS-CoV-2 Spike complexes with the host receptor ACE2 for XBB.1.5+L455F, XBB.1.5+F456L(EG.5) and XBB.1.5+L455F/F456L (FLip) lineages to examine the mechanisms underlying the role of convergent evolution hotspots in balancing ACE2 binding and antibody evasion. Using the ensemble-based mutational scanning of the spike protein residues and physics-based rigorous computations of binding affinities, we identified binding energy hotspots and characterized molecular basis underlying epistatic couplings between convergent mutational hotspots. Consistent with the experiments, the results revealed the mediating role of Q493 hotspot in synchronization of epistatic couplings between L455F and F456L mutations providing a quantitative insight into the mechanism underlying differences between XBB lineages. Mutational profiling is combined with network-based model of epistatic couplings showing that the Q493, L455 and F456 sites mediate stable communities at the binding interface with ACE2 and can serve as stable mediators of non-additive couplings. Structure-based mutational analysis of Spike protein binding with the class 1 antibodies quantified the critical role of F456L and F486P mutations in eliciting strong immune evasion response. The results of this analysis support a mechanism in which the emergence of EG.5 and FLip variants may have been dictated by leveraging strong epistatic effects between several convergent revolutionary hotspots that provide synergy between the improved ACE2 binding and broad neutralization resistance. This interpretation is consistent with the notion that functionally balanced substitutions which simultaneously optimize immune evasion and high ACE2 affinity may continue to emerge through lineages with beneficial pair or triplet combinations of RBD mutations involving mediators of epistatic couplings and sites in highly adaptable RBD regions.

## Introduction

The extensive body of structural and biochemical studies conducted on the Spike (S) glycoprotein of the SARS-CoV-2 virus has yielded pivotal insights into the mechanisms governing virus transmission and immune evasion. This glycoprotein, responsible for viral entry into host cells, undergoes notable conformational changes between closed and open states that are orchestrated by the flexible amino (N)-terminal S1 subunit, comprising the N-terminal domain (NTD), the receptor-binding domain (RBD), and two structurally conserved subdomains—SD1 and SD2.^1–9^ The dynamic interplay of these structural domains is essential for modulating conformational transitions in the S protein between the RBD-down closed state and the RBD-up open state which in turn facilitates a spectrum of functional responses. Moreover, functional motions of the NTD, RBD, and SD1 and SD2 subdomains are synchronized as the S1 subunit engages in global movements that are coordinated through long-range communications with the structurally rigid S2 subunit.^10–15^ The cooperative interplay of the S1 and S2 intrinsic dynamics and functional motions plays a pivotal role in mediating essential interactions between the S protein and the host cell receptor ACE2, as well as governing a broad spectrum of the S interactions with distinct classes of antibodies, influencing the host immune responses triggered by the virus. The ability of the S protein to navigate between distinct structural states is fundamental for its efficacy in recognizing and binding to host cell receptors and evading immune detection. The wealth of information derived from biophysical studies has significantly enhanced our understanding of the S protein trimer shedding light on the intricate interplay of thermodynamics and kinetics governing spike mechanisms. These studies have unveiled a nuanced picture of how mutations and long-range interactions between the dynamic S1 subunit and a more rigid S2 subunit drive coordinated structural alterations within the S protein trimer which determine the dynamic equilibrium and affect population shifts between the open and closed RBD which are central to modulating interactions with various binding partners, virus transmission and immune responses.^16–18^ The growing wealth of cryo-electron microscopy (cryo-EM) and X-ray structures for the S protein variants of concern (VOCs) has significantly enriched our understanding of the evolutionary adaptability of the S protein and diversity of molecular mechanisms. These structural studies revealed multiple functional states of the S protein and interactions with antibodies leading to a complex dynamic landscape and a diverse array of binding epitopes that ultimately determine the virus fitness and balance of between host receptor binding and immune escape.^19–28^ The recent cryo-EM structures and biochemical analyses of the S trimers for various subvariants of the Omicron variant, including BA.1, BA.2, BA.3, and BA.4/BA.5 demonstrated a noteworthy decrease in the binding affinity for the BA.4/BA.5 subvariants, confirming higher binding affinities for BA.2 compared to other Omicron variants.^29,30^ Structural and biophysical investigations of the Omicron BA.2.75 variant indicated that, at neutral pH, the BA.2.75 S-trimer displayed the highest thermal stability among the Omicron variants followed by BA.1, BA.2.12.1, BA.5, and BA.2 variants. Surface plasmon resonance (SPR) experiments conducted for multiple Omicron subvariants, including BA.1, BA.2, BA.3, BA.4/5, BA.2.12.1, and BA.2.75, revealed that the BA.2.75 subvariant exhibited a notably higher ACE2 binding affinity that was approximately 4-6 times greater than the other Omicron variants, further underlining the distinctive features of the BA.2.75 subvariant.^31–33^ The emergence of the XBB.1 subvariant within the Omicron lineage provided a compelling example of viral evolution, representing a descendant of BA.2 and exhibiting a recombinant nature with BA.2.10.1 and BA.2.75 sublineages. A closely related variant, XBB.1.5, shares a striking similarity with XBB.1 and is characterized by a single rare two-nucleotide substitution in the RBD compared to the ancestral strain.^34^ Biophysical studies investigating the binding of the S trimer with ACE2 for Omicron variants, including BA.2, BA.4/5, BQ.1, BQ.1.1, XBB, and XBB.1, revealed that the binding affinities of BQ.1 and BQ.1.1 were comparable to that of the BA.4/5 spike, while XBB and XBB.1 exhibited binding affinities similar to that of the BA.2 variant. Notably, this study reported alarming antibody evasion by Omicron BQ.1, BQ.1.1, XBB, and XBB.1 variants, indicating that monoclonal antibodies effective against the original Omicron variant were largely inactive against these new subvariants.^35^ In particular, XBB.1.5 variant demonstrated immune evasion capabilities comparable to XBB.1 but displayed a potential growth advantage due to a higher ACE2 binding affinity attributed to a single F486P mutation as compared to the F486S substitution in the XBB.1 variant.^35^ Biochemical studies confirmed the exceptional binding affinity of the XBB.1.5 RBD to ACE2 with a dissociation constant comparable to that of the BA.2.75 variant but significantly stronger than that of XBB.1 and BQ.1.1 variants.^36,37^

These studies showed that XBB.1 and XBB.1.5 can exhibit comparable antibody evasion but much greater transmissibility for the XBB.1.5 variant suggesting that for these Omicron lineage the enhanced receptor-binding affinity can confer higher growth advantages.^36^ Moreover, the stronger ACE2 binding of XBB.1.5 could also strengthen its tolerance to further immune escape mutations.^36^ Cryo-EM analysis of the XBB.1.5 S ectodomain confirmed its immune evasion capabilities that are similar to XBB.1 while highlighting a higher ACE2 binding affinity, likely contributing to the growth advantage and increased transmissibility observed in the XBB.1.5 variant.^38^ The immune-evasive XBB lineages continue to evolve accumulating more S mutations, such as R403K, V445S, L455F, F456L and K478R leading to further shift in antigenicity and escape from neutralizing antibodies elicited by repeated vaccination and infection. Since March 2023, sublineages of the XBB variant harboring the F486P substitution (XBB.1.5 and XBB.1.9) dominated worldwide (https://nextstrain.org/ncov/gisaid/global/6m).^39^ Recently, XBB.1.16 sublineage emerged in various countries including the US. As of October 2023, XBB sublineages XBB.1.5 and XBB.1.16, sharing F486P substitution have become predominant worldwide (https://nextstrain.org/)^39^ XBB.1.16 emerged independently from XBB.1.5 and compared to XBB.1.5, XBB.1.16 has two substitutions E180V in the NTD and T478R in the RBD.^40^ Compared to XBB.1.5, XBB.1.16 has E180V and T478R mutations showing a greater growth advantage in the human population as compared to XBB.1 and XBB.1.5 variants, while exhibiting similar immune evasion potential.^40^ The emerging variants display the increased infectivity and transmissibility over previous Omicron variants, and some RBD residues (R346, K356, K444, V445, G446, N450, L452, N460, F486, F490, R493 and S494) are mutated in at least five new independent Omicron sublineages. The waive of XBB descendants, including EG.5 and EG.5.1 (XBB.1.9.2.5.1) that bear additional mutation F456L became one of the currently predominant lineage circulating worldwide.^41^ EG.5 evolved from Omicron XBB.1.9 and harbors only one additional F456L substitution relative to XBB.1.5 while its immediate descendant EG.5.1 features Q52H in the NTD and F456L in the RBD.^41^ Biochemical studies showed that ACE2 binding of EG.5.1 RBD is weaker as compared to that of XBB.1.5 but EG.5.1 exhibits significantly greater immune resistance to the humoral immunity where F456L substitution is implicated as an important determinant in the enhanced immunological potential.^41^ EG.5 and EG.5.1 were found to be moderately more resistant to antibody neutralization as compared to XBB.1.5, particularly showing marked immune resistance for class 1 monoclonal antibodies that is largely mediated by single F456L mutation on the RBD.^42^ This observation was also confirmed in studies of immune evasion of the EG.5.1 subvariant showing that F456L mutation rather than Q52H drives the enhanced neutralization escape of EG.5.1 where binding to class 1 monoclonal antibodies is markedly reduced while binding to S309, a class 3 monoclonal antibody is not appreciably affected.^43^ These studies consistently suggested that the epidemic spread of the EG.5.1 lineage may have been primarily due to the increased immune escape.

EG.5.1 further evolved with a descendant lineage harboring L455F (EG.5.1+L455F) and named HK.3 (XBB.1.9.2.5.1.1.3).^44^ XBB subvariants bearing both L455F and F456L combination of flipped substitutions are termed “FLip” variants that continue to grow and are present in > 20% of global XBB in samples collected at the beginning of September 2023, doubling roughly every 3 weeks.^39^ In China, FLip has increased particularly quickly, mostly as HK.3 (EG.5.1.1.3), crossing 50% in early September with a relative doubling time of 10 days. The FLip variants include also JG.3 (XBB.1.9.2.5.1.3.3), JF.1 (XBB.1.16.6.1), GK.3 (XBB.1.5.70.3) and JD.1.1 that emerged convergently indicating that acquisition of L455F/F456L “combo” can confer a growth advantage to XBB in the human population.^44,45^ In XBB FLip lineages, extra spike mutation A475V appears to be beneficial promoting emergence of JD.1.1 (XBB.1.5.102.1.1), GK.3.1 (XBB.1.5.70.3.1), GK.4 (XBB.1.5.70.4), GW.5.1.1 (XBB.1.19.1.5.1.1), and FL.15.1.1 (XBB.1.9.1.15.1.1). JD.1.1 has three additional mutations compared to XBB.1.5, including the FLip mutations L455F and F456L as well as A475V.^45^ Convergent evolution of XBB lineages on the RBD sites L455F and F456L resulted in EG.5, FL.1.5.1, XBB.1.5.70, and HK.3 variants.^46^

Biochemical studies of ACE2 binding with XBB.1.5, XBB.1.5+L455F, XBB.1.5+F456L and XBB.1.5+L455F+F456L by surface plasmon resonance (SPR) assays revealed that L455F reduced ACE2 affinity, while the adjacent residue flipping of L455F and F456L leads to the enhanced ACE2 binding affinity.^46^ Furthermore, mutations L455F, F456L, and L455F/F456L induce the enhanced immune escape to the majority of class 1 monoclonal antibodies (Figure 1).

**Figure 1.**
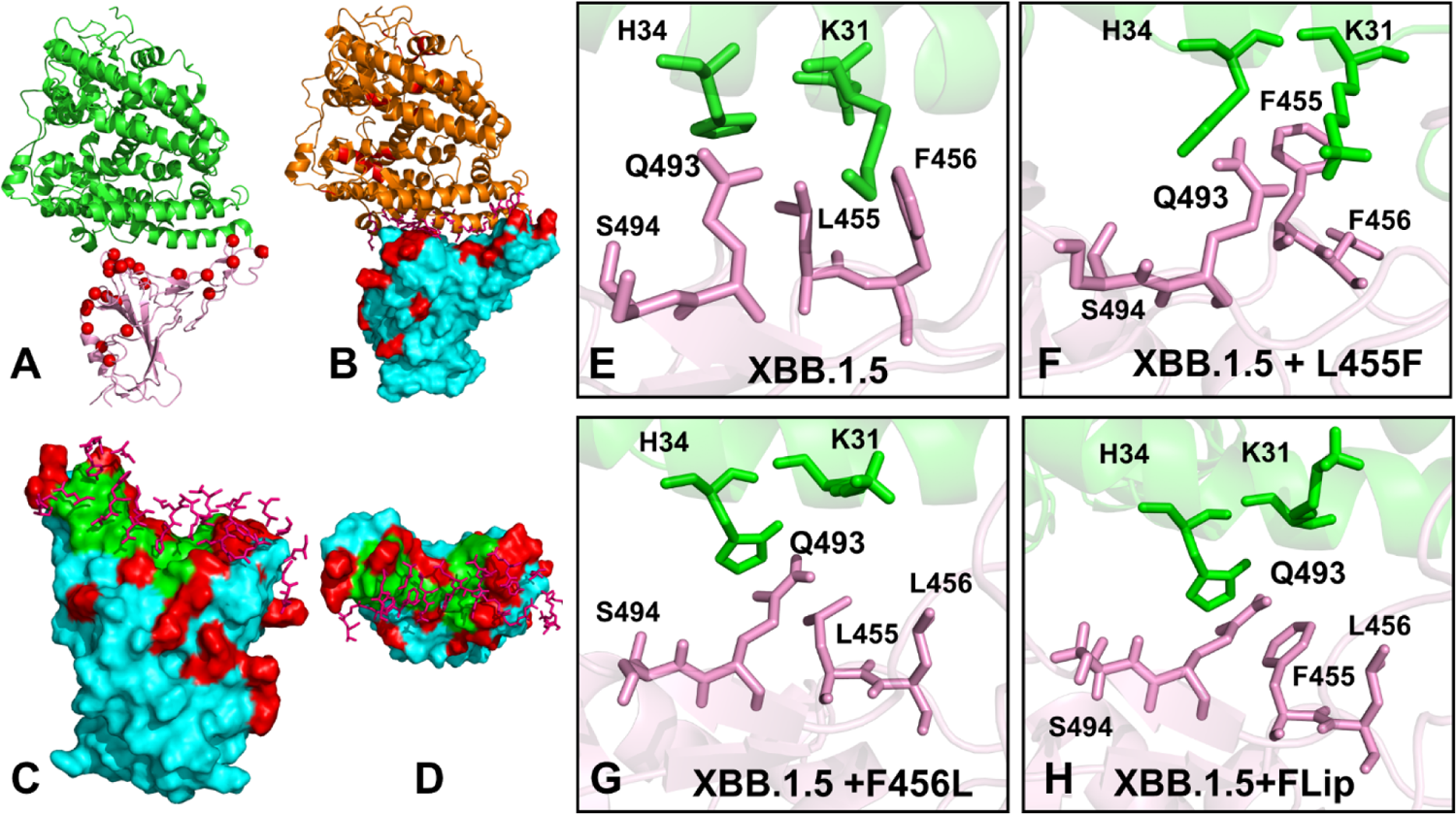
Structural overview of the SARS-CoV-2 S-RBD-ACE2 complexes for Omicron XBB lineages. (A) The cryo-EM structure of the XBB.1.5 RBD-ACE2 complex (pdb id 8WRL). The RBD is shown in pink ribbons, the ACE2 receptor is in green ribbons. The XBB.1.5 RBD mutational sites G339H, R346T, L368I, S371F, S373P, S375F, T376A, D405N, R408S, K417N, N440K, V445P, G446S, N460K, S477N, T478K, E484A, F486P, F490S, R493Q, Q498R, N501Y, Y505H are shown in red spheres. (B) The surface-based representation of XBB.1.5 RBD-ACE2 complex. The RBD is pink surface, ACE2 is in orange ribbons. The XBB.1.5 mutational positions are in red. (C,D) The front and top view of the S-RBD binding epitope for the XBB.1.5 RBD-ACE2 complex. The RBD is in cyan surface, The binding epitope is in green surface, XBB.1/XBB.1.5 Omicron mutations are in red. The ACE2 binding interface residues are shown in sticks. (E) A closeup of the binding interface residues Q493, L455 and F456 in the cryo-EM structure of the XBB.1.5 RBD-ACE2 complex. The RBD residues are in pink sticks, the ACE2 residues H3 and K31 are in green sticks. (F) A closeup of the binding interface residues Q493, L455F and F456 in the AF2-predicted best model of the XBB.1.5+L455F complex with ACE2. (G) A closeup of the binding interface residues Q493, L455 and F456L in the AF2-predicted best model of the XBB.1.5+F456L complex with ACE2. (H) A closeup of the binding interface residues Q493, L455F and F456L in the AF2-predicted best model of the XBB.1.5+ FLip complex with ACE2. The binding interface residues are shown in pink sticks for RBD sites and green sticks for ACE2 sites. The RBD interface residues Q493, L455F and F456L undergo noticeable rearrangements in the XBB.1.5+FLip complex.

The Omicron subvariant BA.2.86 derived from BA.2 variant exhibits significant genetic differences compared to its predecessors.^47–51^ A comparative functional analysis of immune evasion and infectivity for BA.2.86, EG.5.1 and FLip variants showed that BA.2.86 is less resistant to neutralization by bivalent boosted sera compared to XBB.1.5, EG.5.1, and FLip variants.^52^ Notably, FLip variant exhibited stronger immune escape than its parental variant XBB.1.5 due to both L455F and F456L mutations. JN.1 is a variant of BA.2.86 which emerged independently from Omicron BA.2 and harbors an additional L455S mutation responsible for enhanced immune escape.^53^ A comparative biochemical analysis using surface plasmon resonance (SPR) assays showed a notable reduction in ACE2 binding affinity for JN.1 indicating that its enhanced immune evasion capabilities come at the expense of reduced ACE2 binding.^53^ This study also showed that A475V mutation in JD.1.1 (XBB.1.5 + FLip + A475V) can modestly reduce binding affinity while enhancing immune evasion compared to HK.3 (XBB.1.5 + FLip). Interestingly, with single additional L455S mutation compared to its predecessor BA.2.86 and resulting increased resistance to humoral immunity, the JN.1 variant rapidly became predominant in Europe surpassing both BA.2.86 and the “FLip” (L455F+F456L) strains.^53^ The evolutionary pattern of XBB.1.5, E.5.1. and FLip variants suggested a predominant role of immune evasion during virus adaptation where compensatory mutations L455GF/F456L may have emerged in combination to restore the decreased ACE2 affinity. At the same time, evolutionary path of BA.2 derived BA.2.86 and JN.1 variants that incorporated L455S mutation highlights the importance of variants featuring high ACE2 binding affinity and distinct antigenicity, where initially weaker immune evasion can be bolstered by mutating L455 and F456 positions.

Deep mutational scanning (DMS) experiments and functional studies suggested that evolutionary windows for the Omicron variants could be enhanced through epistatic interactions between variant mutations and the broader mutational repertoire.^54–56^ Epistasis is a genetic phenomenon that the effect of one mutation can be altered depending on the presence of other mutations, resulting in non-additive impacts of mutations on specific functions. DMS experiments and systematic mapping of the epistatic interactions between the Omicron RBD mutations showed evidence of compensatory epistasis in which immune escape mutations can individually reduce ACE2 binding but are compensated through epistatic couplings with affinity-enhancing Q498R and N501Y mutations.^57,58^ Recent evolutionary studies revealed strong epistasis between pre-existing substitutions in BA.1/BA.2 variants and antibody resistance mutations acquired during selection experiments, suggesting that epistasis can also lower the genetic barrier for antibody escape.^59^ DMS analysis of the complete XBB.1.5 and BA.2 S proteins measured the effect of over 9,000 mutations on ACE2 binding, cell entry, or immune escape^60^ revealing that mutations outside the RBD ca impact ACE2 binding and that strongest immune escape mutations to the XBB.1.5 are in the RBD sites 357, 371, 420, the 440-447 loop, 455-456, and 473. While many neutralization escape mutations that are strongly deleterious to ACE2 binding (such as mutations at Y473) are not common among circulating variants, several DMS-confirmed escape mutations that are tolerant to ACE2 binding emerged in recent XBB and BA.2 variants, including F456 mutations in EG.5.1 and many other XBB variants and L455 mutations in HK.3.1 and JN.1 variants.^60^ Another study examined the preference of each RBD mutation on antibody escape, human ACE2 binding, and RBD stability using BA.5-based DMS profiles and neutralization profiles against XBB.1.5 to identify the evolutionary trends of the XBB.1.5 lineage revealing the most important functional hotspots at R403S/K, N405K, N417Y, Y453S/C/F, L455W/F/S, F456C/V/L and H505Y/D.^61^

Recent DMS experiments examined the impacts of all mutational changes and single-codon deletions in the BQ.1.1 and XBB.1.5 RBDs on ACE2-binding affinity and RBD folding efficiency, revealing the expanded character of epistatic couplings between RBD residues in addition to dramatic epistatic perturbations induced by N501Y, namely prominent epistatic interactions between R493Q reversed mutations and mutations at positions Y453, L455, and F456 that define the newly emerging EG.5.1 and FLip lineages.^62^ Strikingly, the epistatic interactions between these sites are background-specific as mutations Y453W and F456L decrease ACE2-binding affinity 2.2- and 6.6-fold in the Omicron BA.2 background but enhance ACE2-binding affinity 7.1- and 1.9-fold in XBB.1.5 background, while L455W enhances ACE2-binding affinity 2.5-fold in BA.2 but decreases affinity 7.4-fold in XBB.1.5 variant.^62^ Convergent evolution of the XBB lineages showed that coordination of evolutionary paths at different sites may be largely due to epistatic, rather than random selection of mutations.^63,64^

Computer simulation studies provided important atomistic insights into understanding the dynamics of the SARS-CoV-2 S protein and the effects of Omicron mutations on conformational plasticity of the S protein states and their complexes with diverse binding partners.^65–70^ Recent microsecond multiple molecular dynamics (MD) simulations examined the kinetic and thermodynamic contributions of VOC’s mutations showing lower thermodynamic stability with higher kinetic fluctuations, compared to S proteins of ancestral strains which explained the higher probability to attain open S form and increased exposure to ACE2 binding.^70^ Conformational dynamics and allosteric modulation of SARS-CoV-2 S in the absence or presence of ligands was studied using smFRET imaging assay, showing presence of long-range allosteric control of the RBD equilibrium, which in turn regulates the exposure of the binding site and antibody binding.^71^ Integrative computational modeling approaches revealed that the S protein could function as an allosteric regulatory machinery controlled by stable allosteric hotspots acting as drives drivers and regulators of spike activity.^72–77^ By combining atomistic simulations and a community-based network model of epistatic couplings we found that convergent Omicron mutations such as G446S (BA.2.75, BA.2.75.2, XBB), F486V (BA.4, BA.5, BQ.1, BQ.1.1), F486S, F490S (XBB.1), F486P (XBB.1.5) can display epistatic relationships with the major stability and binding affinity hotspots which may allow for the observed broad antibody resistance.^75^ Analysis of conformational dynamics, binding and allosteric communications in the Omicron BA.1, BA.2, BA.3 and BA.4/BA.5 spike protein complexes with the ACE2 host receptor characterized regions of epistatic couplings that are centered at the binding affinity hotspots N501Y and Q498R enabling accumulation of multiple Omicron immune escape mutations at other sites.^76^ MD simulations and Markov state models systematically characterized conformational landscapes and identify specific dynamic signatures of the early Omicron variants BA.1, BA.2, BA.3 and BA.4/BA.5 variants and recent highly transmissible XBB.1, XBB.1.5, BQ.1, and BQ.1.1 Omicron variants and their complexes showing that convergent mutation sites could control evolution allosteric pockets through modulation of conformational plasticity in the flexible adaptable regions.^77^ Recent computational studies suggested that Omicron mutations have variant-specific effect on conformational dynamics changes in the S protein including allosterically induced plasticity at the remote regions leading to the formation and evolution of druggable cryptic pockets.^78–80^

In this study, we set out to understand complex effects of convergent mutations in the XBB lineages XBB.1.5, XBB.1.5+L455F, XBB.1.5+F456L (EG.5) and XBB.1.5+L455/F456L (FLip) on balancing dynamic, binding energetics with ACE2 and effective evasion of neutralizing antibodies. The lack of molecular details on structure, dynamics and binding energetics of the EG.5 and FLIp RBD binding with the ACE2 receptor and antibodies provides a considerable challenge that needs to be addressed to rationalize the experimental data and establish the atomistic basis for the proposed molecular mechanisms. To do that we employed AI-enabled integrative computational approach in which structure and conformational ensembles of the of the XBB.1.5, XBB.1.5+L455F, XBB.1.5+F456L, EG.5 and FLip variants and their RBD-ACE2 complexes were first predicted using AlphaFold2 (AF2) methods^81,82^ using shallow multiple sequence alignment (MSA).^83–85^ Using several structural alignment metrics we evaluate and rank structural models and analyze the conformational heterogeneity of the predicted multiple conformations. Microsecond atomistic MD simulations of these systems are then conducted to validate the AI-generated conformational ensembles and identify differences in dynamics signatures of the XBB variants. We also perform an ensemble-based mutational scanning of the RBD residues in the XBB.1.5+L455F, EG.5 and XBB.1.5+L455F/F456L FLip RBD-ACE2 complexes in the XBB.1.5 background to identify binding energy hotspots and characterize epistatic couplings between convergent mutational hotspots. We combined mutational profiling of the RBD residues with Molecular Mechanics/Generalized Born Surface Area (MM-GBSA) approach for binding affinity computations of the RBD-ACE2 complexes across all studied XBB subvariants. By combining mutational profiling of the RBD residues in the predicted complexes with network-based community decomposition analysis and clique-based model of epistatic couplings we explore epistatic interactions in the Omicron RBD-ACE2 complexes. We also conduct mutational scanning of the S RBD complexes with several class 1 monoclonal antibodies to quantify the effect of convergent Omicron mutations on immune escape and the mechanism underlying balance between ACE2 binding and immune evasion.

## Materials and Methods

### AI-based structural modeling and statistical assessment of AF2 models

Structural prediction of the XBB.1, XBB.1.5, XBB.1.5+L455F, XBB.1.5+F456L (EG.5) and FLIp RBD-ACE2 complexes were carried out using AF2 framework^83,84^ within the ColabFold implementation^86^ using a range of MSA depths and other parameters. The default MSAs are subsampled randomly to obtain shallow MSAs containing as few as five sequences. We used *max_msa* field to set two AF2 parameters in the following format: *max_seqs:extra_seqs*. Both of these parameters determine the number of sequences subsampled from the MSA (*max_seqs* sets the number of sequences passed to the row/column attention track and *extra_seqs* the number of sequences additionally processed by the main evoformer stack). The lower values encourage more diverse predictions but increase the number of misfolded models. Similar to previous studies showing how MSA depth adaptations may facilitate conformational sampling,^83–85^ MSA depth was modified by setting the AF2 config.py parameters *max_extra_msa* and *max_msa_clusters* to 32 and 16, respectively. We additionally manipulated the *num_seeds* and the *num_recycles* parameters to produce more diverse outputs. We use *max_msa*: 16:32, *num_seeds*: 4, and *num_recycles*: 12. AF2 makes predictions using 5 models pretrained with different parameters, and consequently with different weights. To generate more data, we set the number of recycles to 12, which produces 14 structures for each model starting from recycle 0 to recycle 12 and generating a final refined structure. Recycling is an iterative refinement process, with each recycled structure getting more precise. Each of the AF2 models generates 14 structures, amounting to 70 structures in total. In addition, we also predicted one more structure using AF2 with the default and ‘auto’ parameters serving as a baseline structure for prediction and variability analysis.

AF2 models were ranked by Local Distance Difference Test (pLDDT) scores (a per-residue estimate of the prediction confidence on a scale from 0 to 100), quantified by the fraction of predicted Cα distances that lie within their expected intervals. The values correspond to the model’s predicted scores based on the lDDT-Cα metric, a local superposition-free score to assess the atomic displacements of the residues in the model.^81,82^ Structural models were compared to the available experimental structure of the XBB.1 RBD-ACE2 (pdb id 8IOV) and just released structure of the XBB.1.5 RBD-ACE2 complex (pb id 8WRL) using structural alignment as implemented in TM-align.^87^ An optimal superposition of the two structures is then built and TM-score is reported as the measure of overall accuracy of prediction for the models. Several other structural alignment metrics were used including the global distance test total score GDT_TS of similarity between protein structures and implemented in the Local-Global Alignment (LGA) program^88^ and the root mean square deviation (RMSD) superposition of backbone atoms (C, Cα, O, and N) calculated using ProFit (http://www.bioinf.org.uk/software/profit/).

### All-Atom Molecular Dynamics Simulations

The structures of the XBB.1 RBD-ACE2 (pdb id 8IOV) and XBB.1.5 RBD-ACE2 (pdb id 8WRL) are obtained from the Protein Data Bank. The best AF2 models from five predicted conformations obtained with AF2 default parameters for each of XBB lineages were respectively selected as starting structure for MD simulations of XBB.1.5, XBB.1.5+L455F, XBB.1.5 +F456L (EG.5), and FLip RBD-ACE2 complexes. For simulated structures, hydrogen atoms and missing residues were initially added and assigned according to the WHATIF program web interface.^89^ The missing regions are reconstructed and optimized using template-based loop prediction approach ArchPRED.^90^ The side chain rotamers were refined and optimized by SCWRL4 tool.^91^

The protonation states for all the titratable residues of the ACE2 and RBD proteins were predicted at pH 7.0 using Propka 3.1 software and web server.^92,93^ The protein structures were then optimized using atomic-level energy minimization with composite physics and knowledge-based force fields implemented in the 3Drefine method.^94,95^ We considered glycans that were resolved in the structures. NAMD 2.13-multicore-CUDA package^96^ with CHARMM36 force field^97^ was employed to perform 1µs all-atom MD simulations for the Omicron RBD-ACE2 complexes. The structures of the SARS-CoV-2 S-RBD complexes were prepared in Visual Molecular Dynamics (VMD 1.9.3)^98^ and with the CHARMM-GUI web server^99,100^ using the Solutions Builder tool. Hydrogen atoms were modeled onto the structures prior to solvation with TIP3P water molecules^101^ in a periodic box that extended 10 Å beyond any protein atom in the system. To neutralize the biological system before the simulation, Na^+^ and Cl^−^ ions were added in physiological concentrations to achieve charge neutrality, and a salt concentration of 150 mM of NaCl was used to mimic a physiological concentration. All Na^+^ and Cl^−^ ions were placed at least 8 Å away from any protein atoms and from each other. MD simulations are typically performed in an aqueous environment in which the number of ions remains fixed for the duration of the simulation, with a minimally neutralizing ion environment or salt pairs to match the macroscopic salt concentration.^102^ All protein systems were subjected to a minimization protocol consisting of two stages. First, minimization was performed for 100,000 steps with all the hydrogen-containing bonds constrained and the protein atoms fixed. In the second stage, minimization was performed for 50,000 steps with all the protein backbone atoms fixed and for an additional 10,000 steps with no fixed atoms. After minimization, the protein systems were equilibrated in steps by gradually increasing the system temperature in steps of 20 K, increasing from 10 K to 310 K, and at each step, a 1ns equilibration was performed, maintaining a restraint of 10 Kcal mol^−1^ Å^−2^ on the protein C_α_ atoms. After the restraints on the protein atoms were removed, the system was equilibrated for an additional 10 ns. Long-range, non-bonded van der Waals interactions were computed using an atom-based cutoff of 12 Å, with the switching function beginning at 10 Å and reaching zero at 14 Å. The SHAKE method was used to constrain all the bonds associated with hydrogen atoms. The simulations were run using a leap-frog integrator with a 2 fs integration time step. The ShakeH algorithm in NAMD was applied for the water molecule constraints. The long-range electrostatic interactions were calculated using the particle mesh Ewald method^103^ with a cut-off of 1.0 nm and a fourth-order (cubic) interpolation. The simulations were performed under an NPT ensemble with a Langevin thermostat and a Nosé–Hoover Langevin piston at 310 K and 1 atm. The damping coefficient (gamma) of the Langevin thermostat was 1/ps. In NAMD, the Nosé–Hoover Langevin piston method is a combination of the Nosé– Hoover constant pressure method^104^ and piston fluctuation control implemented using Langevin dynamics.^105,106^ An NPT production simulation was run on equilibrated structures for 1µs keeping the temperature at 310 K and a constant pressure (1 atm).

### Mutational Scanning and Binding Free Energy Computations

Mutational scanning analysis of the binding epitope residues was conducted for the all studied RBD-ACE2 complexes. Each binding epitope residue was systematically mutated using all substitutions and corresponding binding free energy changes were computed using BeAtMuSiC approach.^107–109^ This approach considers the effect of the mutation on the strength of the interactions at the interface and on the overall stability of the complex. The binding free energy of protein-protein complex can be expressed as the difference in the folding free energy of the complex and folding free energies of the two protein binding partners:

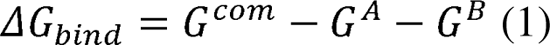

The change of the binding energy due to a mutation was calculated then as the following:

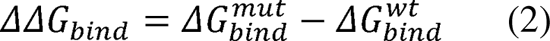

We compute the ensemble-averaged binding free energy changes using equilibrium samples from simulation trajectories. The binding free energy changes were computed by averaging the results over 1,000 equilibrium samples for each of the studied systems. The binding free energies were initially computed for the Omicron RBD-ACE2 complexes using the Molecular Mechanics/Generalized Born Surface Area (MM-GBSA) approach.^110,111^ We also evaluated the decomposition energy to assess the energy contribution of each amino acid during the binding of RBD to ACE2.^112,113^ The binding free energy for the each RBD–ACE2 complex was obtained using:

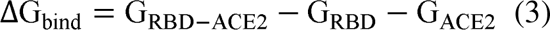

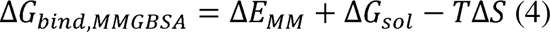

where Δ*E_MM_*is total gas phase energy (sum of Δ*E_internal_*, Δ*E_electrostatic_*, and Δ*Evdw*); Δ*Gsol* is sum of polar (Δ*G_GB_*) and non-polar (Δ*G_SA_*) contributions to solvation.

Here, G_RBD–ACE2_ represent the average over the snapshots of a single trajectory of the MD RBD– ACE2complex, G_RBD_ and G_ACE2_ corresponds to the free energy of RBD and ACE2 protein, respectively. MM-GBSA is employed to predict the binding free energy and decompose the free energy contributions to the binding free energy of a protein–protein complex on per-residue basis as implemented in recent study.^114^ The binding free energy with MM-GBSA was computed by averaging the results of computations over 1,000 samples from the equilibrium ensembles. The entropy contribution was not included in the calculations due to the difficulty of accurately calculating entropy for a large protein–protein complex but equally importantly because the entropic differences between XBB variants for estimates of binding affinities are exceedingly small owing to small mutational changes and preservation of the conformational dynamics.

### Network-Based Community Analysis and Clique-Based Model of Epistatic Interactions

A graph-based representation of protein structures^115,116^ is used to represent residues as network nodes and the inter-residue edges to describe non-covalent residue interactions. The network edges that define residue connectivity are based on non-covalent interactions between residue side-chains. The weights of the network edges in the residue interaction networks are determined by dynamic residue cross-correlations obtained from MD simulations^117^ and coevolutionary couplings between residues measured by the mutual information scores.^118^ The edge lengths in the network are obtained using the generalized correlation coefficients associated with the dynamic correlation and mutual information shared by each pair of residues. The analysis of the interaction networks was done using network parameters such as cliques and communities. The Girvan-Newman algorithm^119^ is used to identify local communities. In this approach, edge centrality (also termed edge betweenness) is defined as the ratio of all the shortest paths passing through a particular edge to the total number of shortest paths in the network. The method employs an iterative elimination of edges with the highest number of the shortest paths that go through them. By eliminating edges, the network breaks down into smaller communities. The algorithm starts with one vertex, calculates edge weights for paths going through that vertex, and then repeats it for every vertex in the graph and sums the weights for every edge. However, in complex and dynamic protein structure networks it is often that number of edges could have the same highest edge betweenness. An improvement of the Girvan-Newman method was implemented, and the algorithmic details of this modified scheme were given in our recent studies.^120^

The k -cliques are complete sub graphs of size k in which each node is connected to every other node. A k -clique community is determined by the Clique Percolation Method^121^ as a subgraph containing k -cliques that can be reached from each other through a series of adjacent *k*-cliques. We have used a community definition according to which in a k -clique community two k - cliques share k-1 or k-2 nodes. Computation of the network parameters was performed using the Clique Percolation Method as implemented in the CFinder program.^122^ Given the chosen interaction cutoff !_min_ we typically obtain communities formed as a union of k = 3 and k = 4 cliques. We assume that residues that belong to the same clique during simulations would have stronger dynamic and energetic couplings leading to synchronization and potentially epistatic effects. To examine the epistatic effect of a mutational site, we compared changes in the k -clique community distributions induced by single and double mutations and calculated the probability by which the two mutational sites belong to the same interfacial 3-clique.^123^

We computed the proportion *P_ab_* of snapshots in the ensemble in which the two mutational sites (*a*, *b*) belong to the interfacial 3-clique:

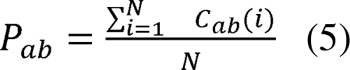

C_ab_(i) = 1 if (*a*,*b*) belong to the same 3-clique. *P_ab_* measures the probability that two sites (*a*, *b*) are kept in some 3-clique due to either direct or indirect interactions. The closer *P_ab_* is to 1, the more likely *a* and *b* tend to have a tight connection and potential local epistasis. To further investigate the effect of mutations on the 3-clique probability, we compared changes in *P_ab_* after single and double mutations. If double mutations have a greater effect on *P_ab_*than single mutations, there may be an epistatic effect between the two sites.

## Results

### AF2-Based Modeling and Prediction of the XBB RBD-ACE2 Complexes and Conformational Ensembles

We proceeded with AF2 structural predictions of the RBD-ACE2 complexes for XBB variants (Figure 1) using AF2 default settings within the ColabFold.^86^ To validate this approach, we first used the predicted best five models for the XBB.1 RBD-ACE2 complex and compared them with the structures of XBB.1 (pdb id 8IOV) and XBB.1.5 complexes (pdb id 8WRL) (Supporting Information, Figures S1, S2). The statistical analysis and confidence assessment metrics of the best five AF2 predictions for XBB.1 and XBB.1.5 RBD-ACE2 complexes are shown as the heatmaps of the MSA sequence coverage and residue-based pLDDT scores estimating the prediction confidence (Supporting Information, Figures S1, S2). The pLDDT scores for the majority of the RBD residues were within ∼ 70-90 range, signaling a fairly high confidence of the predictions, while the lower pLDDT values ∼55-70 were seen for the RBD loops 381-394 and RBM tip region 475-487 reflecting the intrinsic flexibility of these RBD segments. Noticeably, we observed a small divergence in the pLDDT profiles for the best five models of the XBB.1 RBD-ACE2 complex (Supporting Information, Figure S1B) indicating a remarkably high degree of similarity between the best predicted conformations. Some divergence between the models was seen only in the flexible RBM tip (residues 475-487). This is also reflected in the heat maps of the predicted alignment error (PAE) between each residue in the model (Supporting Information, Figure S1C), highlighting the higher confidence of the RBM tip in the best model and moderate decrease in the confidence level for this region in the other models. A similarly high prediction confidence was also observed for the XBB.1.5 RBD-ACE2 complex (Supporting Information, Figure S2). However, we noticed a greater dispersion between the five models in the pLDDT profiles (Supporting Information, Figure S2B) and PAE heatmaps (Supporting Information, Figure S2C) for the flexible RBD regions 381-394, 440-452 and 470-491.

Interestingly, we also found that pLDDT values and PAEs for the RBM tip residues in the best three models (Supporting Information, Figure S2B,C) are higher for XBB.1.5 variant than for the XBB.1 variant. These observations suggested the greater stability of the RBM tip in XBB1.5 which is consistent with the notion that S486P mutation in XBB.1.5 may reduce the loop variation and improve the interactions with the ACE2 receptor. Structural alignment of the best AF2 model with the experimental structure for both XBB.1 and XBB.1.5 yielded the root mean square deviation, RMSD < 0.5 Å (Supporting Information, Figure S1D, S2D). Based on this analysis, we proceeded with AF2 predictions for the XBB.1.5+L455F, XBB.1.5+F456L (EG.5) and FLip RBD-ACE2 complexes (Figure 2). The pLDDT distributions revealed small deviations between the profiles of the best five models for XBB.1.5+L455F (Figure 2A) and progressively increased dispersion between the AF2 models for EG.5 (Figure 2B) and FLip RBD-ACE2 complexes (Figure 2C). The lowered pLDDT values correspond to the RBD regions (residues 450-490) indicating that introduction of F456L and L455F/F456L mutations may moderately increase flexibility of these residues, particularly the RBM tip loop. The inspection of PAE heatmaps showed that the best AF2 model for each of these variants can predict the RBD conformation for all regions with high confidence showing only minor reduction in accuracy for the RBM tip (Figure 2D-F).

**Figure 2.**
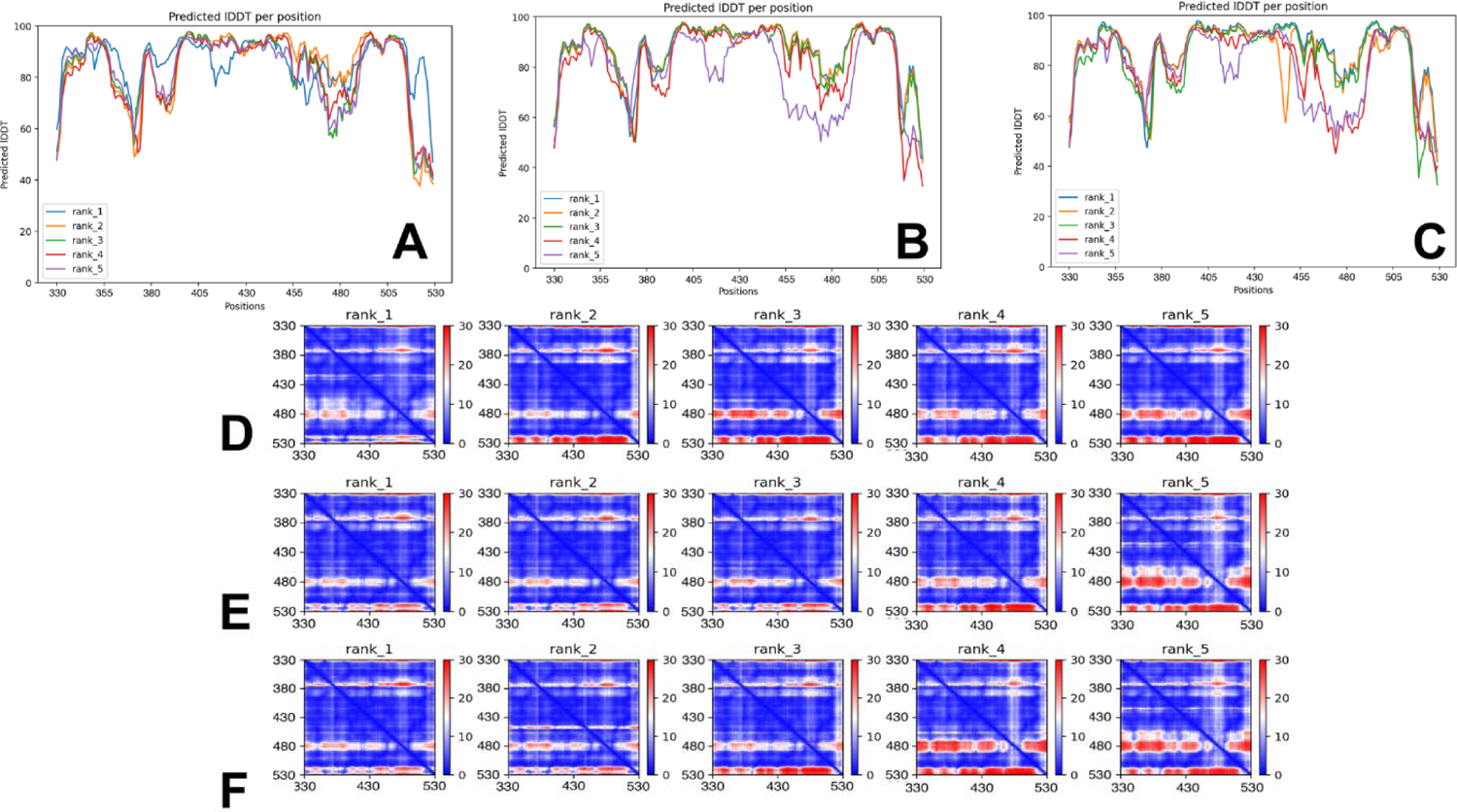
The statistical analysis of AF2 predictions for the XB RBD-ACE2 complexes. The residue-based pLDDT profile for the top five ranked models that are obtained from AF2 predictions of the XBB.1.5+L455F (A), XBB.1.5+F456L (B) and XBB.1.5+L455F/F456L Flip complexes (C). The predicted alignment error (PAE) heatmaps for the top five models for the XBB.1.5+L455F (D), XBB.1.5+F456L (E) and XBB.1.5+L455F/F456L FLip complexes (F). The heat maps are provided for each top ranked model and show the PAE between each residue in the model. The color scale contains three colors to highlight the contrast between the high confidence regions and the low confidence regions.

Structural alignment of the best AF2 model for XBB.1.5+L455F (Figure 3A), XBB.1.5+F456L EG.5 (Figure 3B) and FLip RBD conformations (Figure 3C) with the cryo-EM conformation of XBB.1.5 showed almost perfect overlap and only minor displacements of the RBM tip while preserving the experimentally observed “hook” state (Figure 3). Previous studies showed that the RBM tip may also adopt a highly dynamic “disordered” state in which the tip samples a variety of conformations which is related to the increased variability of the RBD.^124^

**Figure 3.**
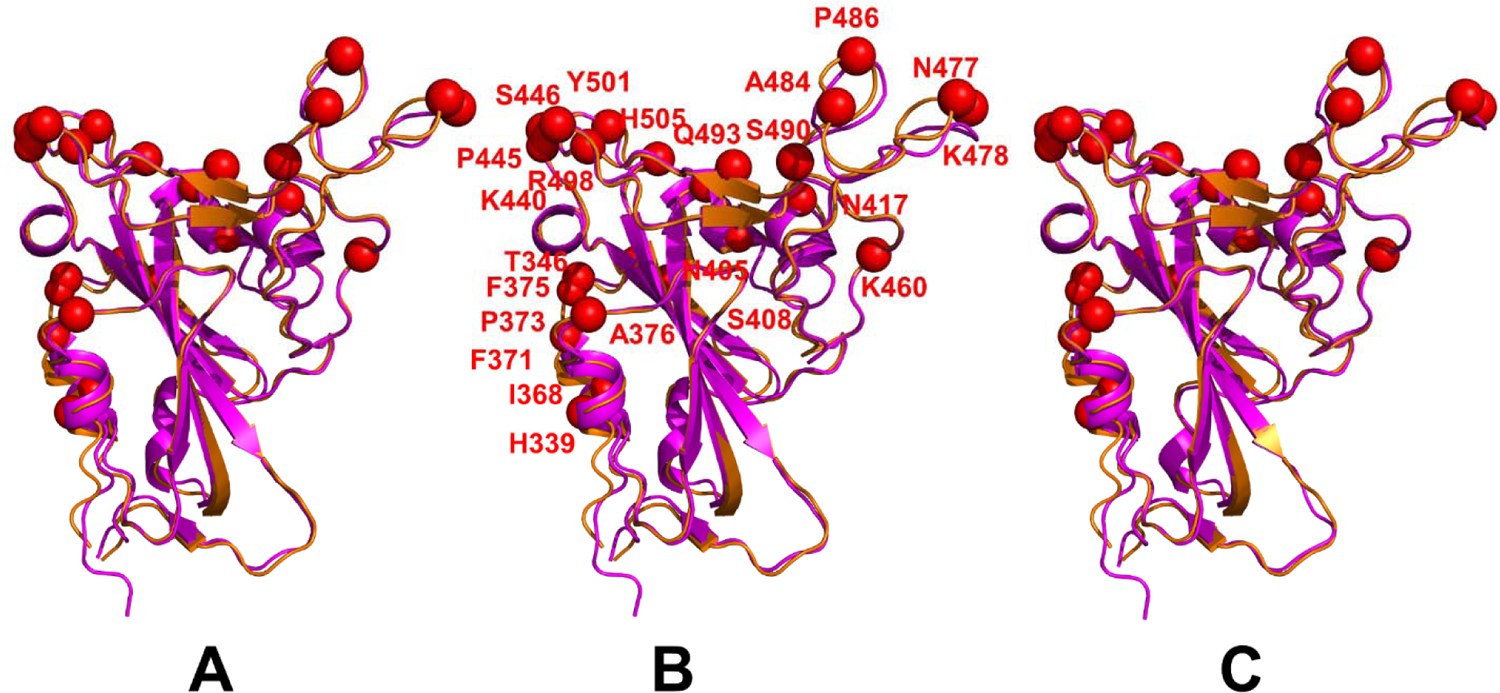
Structural alignment of the best ranked AF2 model with the cryo-EM conformation of XBB.1.5 RBD (pd id 8WRL) for XBB.1.5+L455F ( A), XBB.1.5+F456L EG.5 (B) and XBB.1.5+L455F/F456L RBD conformations (C). The XBB.1.5 RBD conformation (pdb id 8WRL) is shown in orange ribbons. The AF2 predicted XBB RBD models are shown in magenta ribbons. The sites of XBB.1.5 RBD mutations and positions of L455F and F456L mutations are shown in red spheres.

We then proceeded using AF2 methodology with varied MSA depth to predict structural ensembles of the RBD-ACE2 complexes for XBB.1, XBB.1.5, XBB.1.5+L455F, EG.5 and FLip variants. By using two AF2 parameters in the following format: *max_seqs:extra_seqs* we can manipulate the number of sequences subsampled from the MSA (*max_seqs* sets the number of sequences passed to the row/column attention track and *extra_seqs* the number of sequences additionally processed by the main evoformer stack). More diverse predictions are generally encouraged with the lower values and MSA depth adaptations may facilitate conformational sampling^83–85^ Consistent with previous studies, we found changing *max_seq* and *extra_seq* allows to obtain robust predictions with a *max_seq*:*extra_seq* ratio of 256:512 leading to the most diversity of the RBD loops and allowing for reconstruction of the the experimental structure (Figure 4). Using this prediction as reference, we tested several combinations of parameters to employ AF2 with varied MSA depth. Te objective of this analysis was to adequately describe structural ensembles of the RBD-ACE2 complexes in which the experimental structure is the most frequent prediction within the ensemble providing the highest degree of confidence based on pLDDT metric. The ensemble of the predicted AF2 models was evaluated by the pLDDT metric showing strong and consistent peaks at pLDDT ∼85 for all studied XBB variants (Figure 4). In particular, the pLDDT assessments of the structural ensembles for XBB.1 (Figure 4A) and XBB.1.5 complexes (Figure 4B) reveled that the dominant fraction of the ensemble (pLDDT ∼80-85) conforms very closely with the experimental structure, while a minor fraction of the ensemble samples pLDDT values ∼60-80 associated with the native RBD conformations but featuring the increased mobility of the RBD flexible loops For the XBB.1.5 variant, in addition to the major peak of pLDDT ∼85, we observed two additional peaks corresponding to pLDDT ∼75 an pLDDT ∼65 (Figure 4B) which may be interpreted as the increased heterogeneity of the XBB.1.5 ensemble.

**Figure 4.**
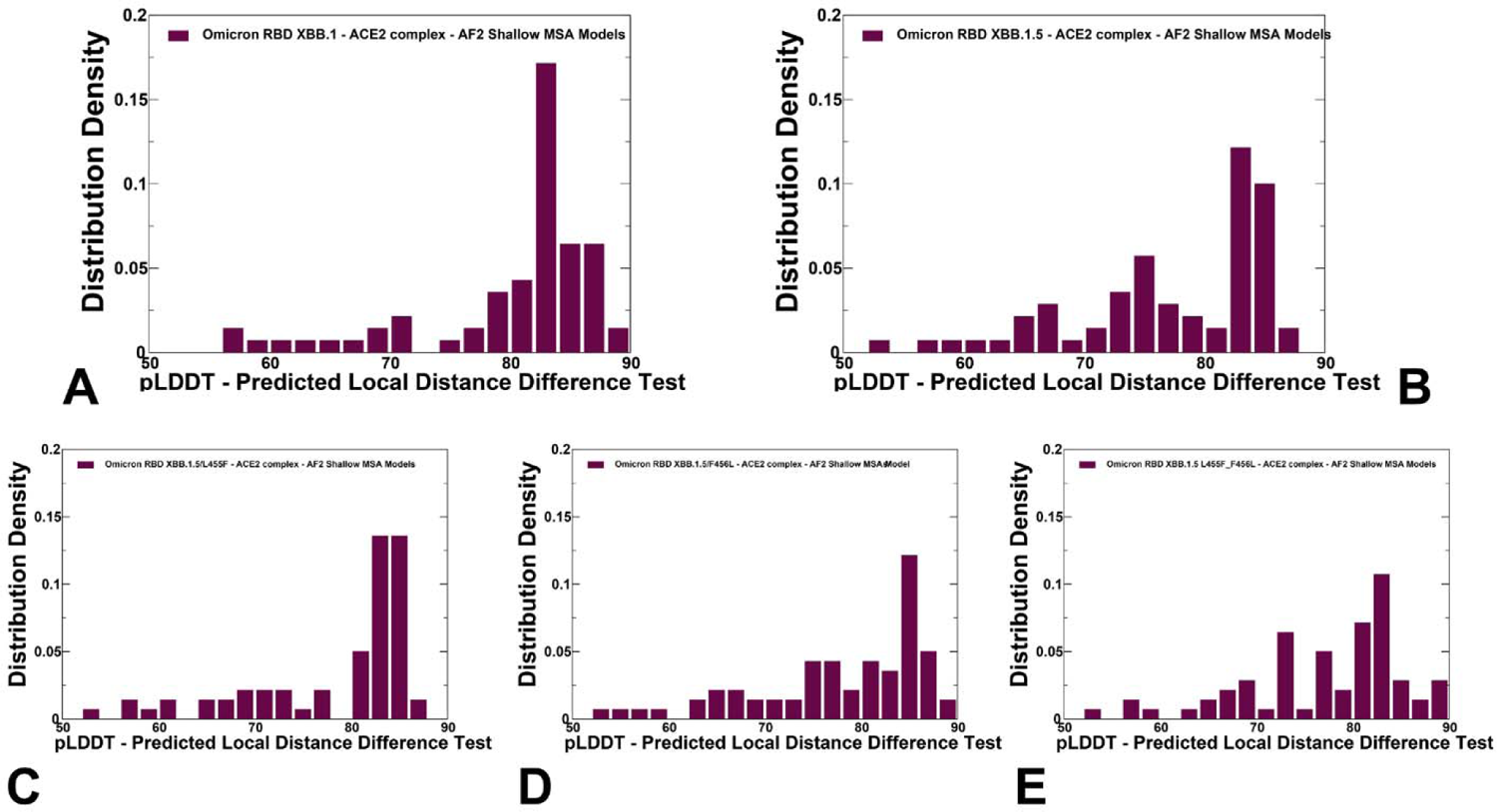
The distributions of the pLDDT metric for the XBB RBD-ACE2 conformational ensembles obtained from AF2-MSA depth predictions. The density distribution of the pLDDT structural model estimate of the prediction confidence for XBB.1 (A) XBB.1.5 (B), XBB.1.5+L455F(C), XBB.1.5+F456L (D) and XBB.1.5+L455F/F456L FLip variants (E).

It is worth noting that that AF2 predictions with pLDDT values ∼70-90 are typically associated with high confidence, the regions with pLDDT values ∼ 50-70 have somewhat lower confidence, while pLDDT < 50 may be a reasonably strong predictor of disorder.^81–84^ Hence, the obtained conformational ensembles represent high confidence structural predictions that can identify and reproduce mobility of the intrinsically flexible RBD regions.

We compared the distributions of the pLDDT values for the XBB.1.5+L455F, EG.5 and FLip RBD-ACE2 complexes (Figure 4C-E) that displayed similar pLDDT distributions featuring major pLDDT peak ∼ 80-90 and indicating high confidence of predictions for conformations in the ensemble. At the same time, we noticed minor but potentially relevant differences, particularly a more dominant major peak pLDDT ∼ 85 for XBB.1.5+L455F and the reduced contribution of alternative peaks, which may indicate a more rigid ensemble of conformations for this subvariant (Figure 4C). According to the pLDDT profiles, conformational heterogeneity may gradually increase in the XBB.1.5+F456L variant and FLip variant (Figure 4E) where the distribution becomes noticeably broader featuring several additional peaks at pLDDT ∼65-75 range. Hence, the AF2 predicted conformational ensemble for the FLip variant displayed greater heterogeneity, particularly in the flexible RBD loops. Using several structural similarity metrics TM-score (Figure 5), GDT TS (Supporting Information, Figure S3) and RMSD (Supporting Information, Figure S4) the prediction accuracy of AF2-MSA depth models was evaluated. The density distributions of TM-scores for XBB.1 (Figure 5A) and XBB.1.5 RBD-ACE2 complexes (Figure 5B) highlighted a considerable similarity between the predicted conformations and the experimental structures with the major peaks corresponding to TM-score ∼0.95. Several minor peaks at TM-score ∼0.7-0.8 seen in the XBB.1.5 variant reflected a moderate variability in the flexible regions. The TM-score density distributions for XBB.1.5+L455F, XBB.1.5+F456L and FLIp conformations (Figure 5C-E) were fairly similar to the XBB.1.5. However, one could notice a slightly more skewed distribution towards TM-score ∼0.9-0.95 for XBB.1.5+L455F suggestive of only minor motions in the flexible regions (Figure 5C). The introduction of F456L produced a marginally increased variability (Figure 5D), while the XBB FLip variant featured the distribution indicative of the increased heterogeneity manifested in mobility of the RBM loops (Figure 5E). Other structural similarity metrics such as the global distance test total score GDT_TS (Supporting Information, Figure 3) and RMSD (Supporting Information, Figure 4) produced distributions indictive of only moderate variability of the RBD conformations in the AF2 structural ensembles across all studied XBB lineages, The RMSD distribution showed that most of the predicted conformations are similar to the cryo-EM structure with the main peak corresponding to RMSD ∼ 0.75-0..85 Å from the experimental structure (Supporting Information, Figure 4).

**Figure 5.**
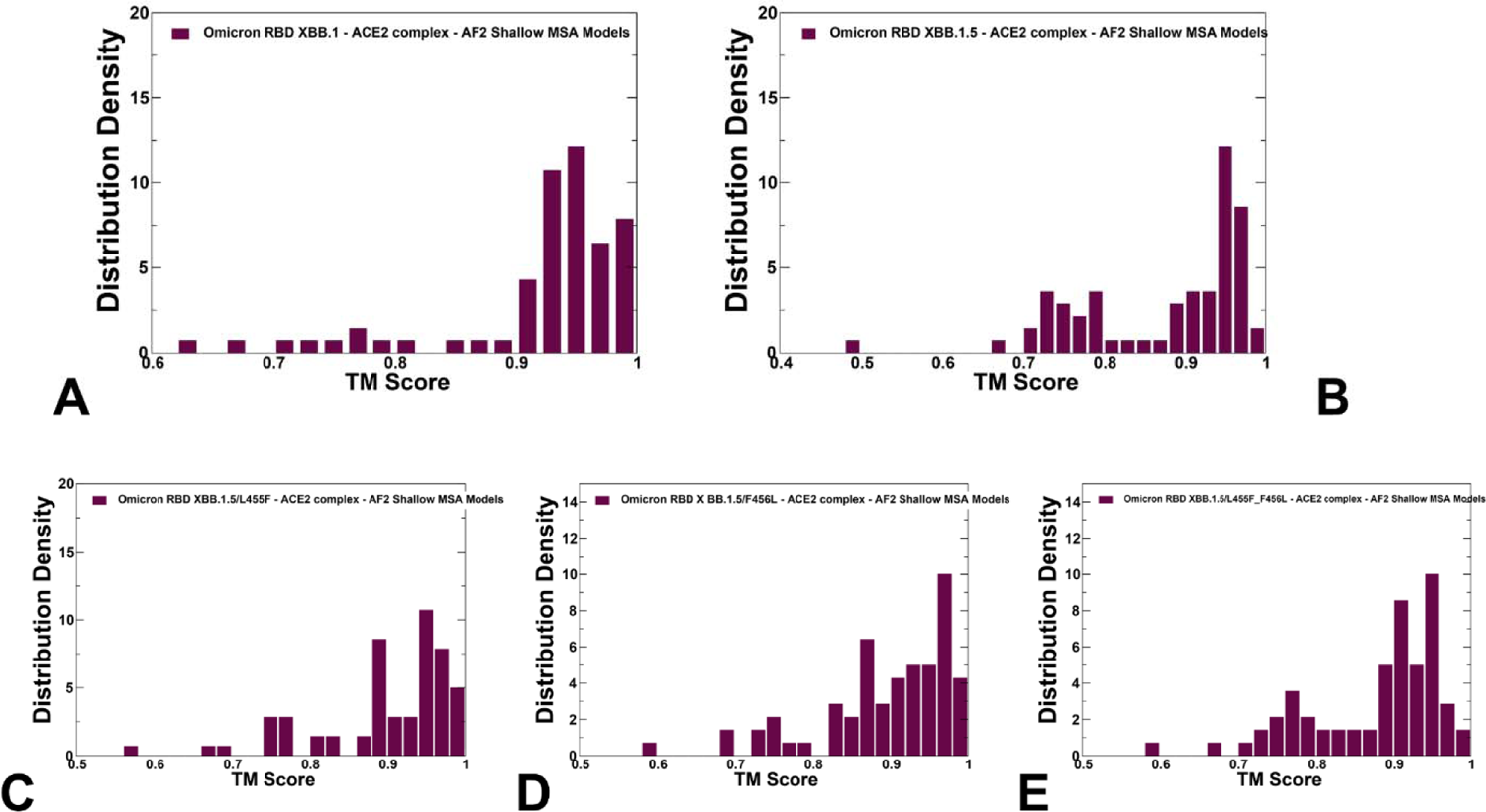
The density distribution of TM-score measuring structural similarity of the predicted RBD conformational ensembles with respect to the structure of the XBB.1.5 RBD-ACE2 (pdb id 8WRL) for XBB.1 (A) XBB.1.5 (B), XBB.1.5+L455F(C), XBB.1.5+F456L (D) and XBB.1.5+L455F/F456L FLip variants (E).

Structural alignment of the AF2 conformational ensembles with the cryo-EM structures illustrated the predicted patterns of RBD mobility in which the RBD core and most of the loops largely remain in their native positions, while most of variability is observed in the RBD residues 470-490 that include important mutational positions S477N, T478K, E484A, F486P, F490S (Figure 6). Consistent with previous MD simulation studies^77^, we observed that the intrinsically flexible RBM tip is maintained in a folded conformation that can be described as a “hook-like” folded motif^124^ that is similar to the experimental conformations and the RBM variability can be attributed to moderate “up” and “down” displacements (Figure 6). The RBD variability and displacements of the RBM tip motif are curtailed in the XBB.1.5/L455F conformation (Figure 6B), while these fluctuations are more pronounced in the XBB.1.5+F456L (Figure 6C) and FLip ensembles (Figure 6D) and are similar to the dynamic changes observed for the XBB.1.5 RBD (Figure 6A). Structural projection of the Omicron XBB.1.5 mutations showed that L455F and F456L substitutions do not induce appreciable dynamic changes in the site of mutations but may moderately affect the mobility of the RBM tip, which suggests a potential for synergistic effects between L455F, F456L and Omicron RBM sites. It can be noticed that the predicted RBD ensembles displayed some variability and dynamic changes around positions E484A, F486P, S477N, T478K (Figure 6). It could also be seen that F486P may experience some fluctuations which is consistent with the presence of many conserved mutations (F486V, F486I, F486S, F486P) seen in other variants making this position a convergent evolutionary hotspot. In general, these results showed that AF2-generated conformational ensembles can accurately reproduce the experimental structures and capture conformational details of the RBD fold and variant-specific functional adjustments of the RBM loop impacting the level of exposure for mutational positions in this region. Importantly, pLDDT assessment of the AF2 models can be used to quantify the variability and extent of dynamic changes in the flexible RBD regions. Based on these findings, we argue that the AF2-predicted conformational plasticity in the RBM region may be functionally significant and reflect the tendency of latest immune-evasive XBB lineages to evolve additional mutations including A475V and K478R from the RBM tip that result in further shift in antigenicity and immune escape from neutralizing antibodies elicited by repeated vaccination.

**Figure 6.**
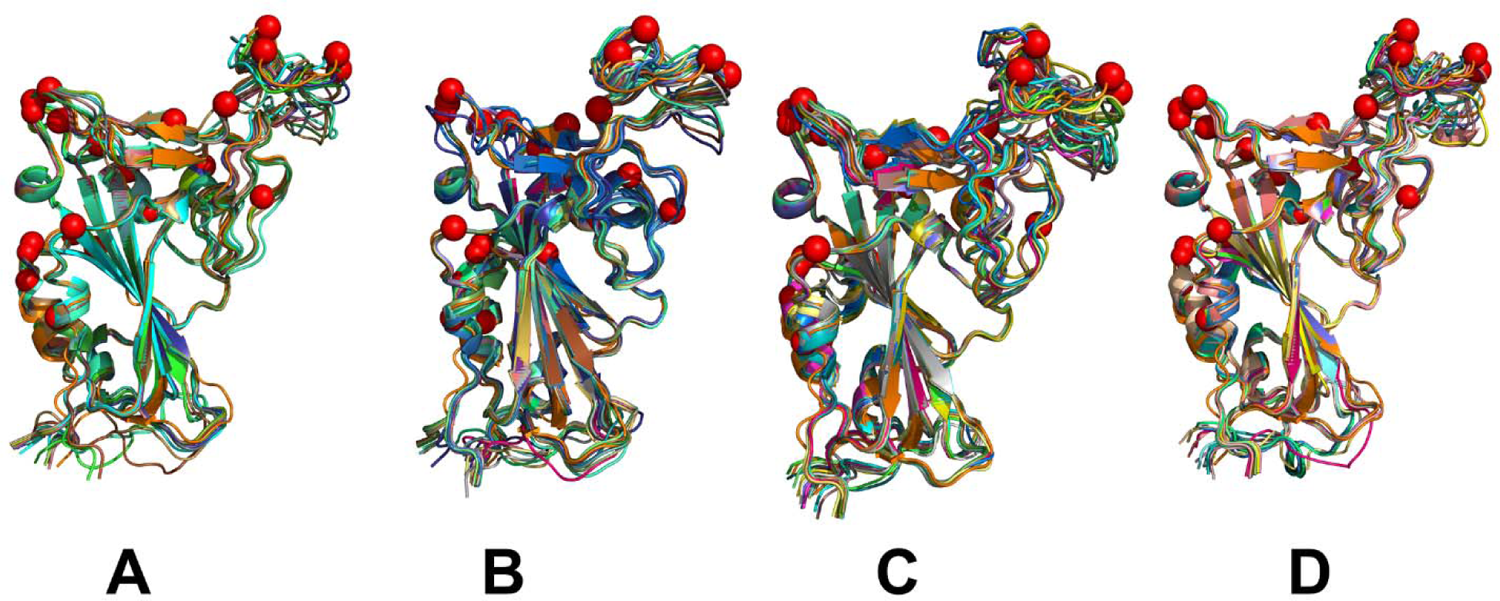
Structural alignment of the AF2-predicted RBD conformational ensembles with high pLDDT values and the cryo-EM structure of the XBB.1.5 RBD-ACE2 complex for XBB.1.5 *(A), XBB.1.5+L455F (B), XBB.1.5+F456l (C) and XBB.1.5+L455F/F456L (D). The RBD conformations are shown in ribbons and the positions of the XBB.1.5 mutational sites G339H, R346T, L368I, S371F, S373P, S375F, T376A, D405N, R408S, K417N, N440K, V445P, G446S, N460K, S477N, T478K, E484A, F486P, F490S, R493Q reversal, Q498R, N501Y, Y505H as well as L455F and F456l are shown in red spheres.

The growth of the XBB variants that combine F456L with A475V or K478R, another mutation known from XBB.1.16 appeared to be beneficial for virus evolution.^39^ According to our observations, in the XBB variants, the mutational sites from the RBM tip undergo moderate displacements around an ordered “hook-like” conformation (Figure 6). It is possible that additional substitutions in these exposed A475 and K478 positions may alter/reduce the fraction of ordered RBM conformations and promote more dynamic RBM ensemble in which the tip samples a variety of more disordered conformations, which is in turn may become beneficial for variant-specific modulation and enhancement of immune escape.

### Atomistic MD Simulations of the XBB RBD-ACE2 ACE2 Complexes

To characterize conformational landscapes and dynamic signatures of the RBD-ACE2 variants for the XBB variants we conducted multiple independent microsecond MD simulations for each of the studied RBD-ACE2 complexes (Table 2, Figure 7) MD simulations revealed important features of the intrinsic conformational dynamics of the RBDs among Omicron variants.

**Figure 7.**
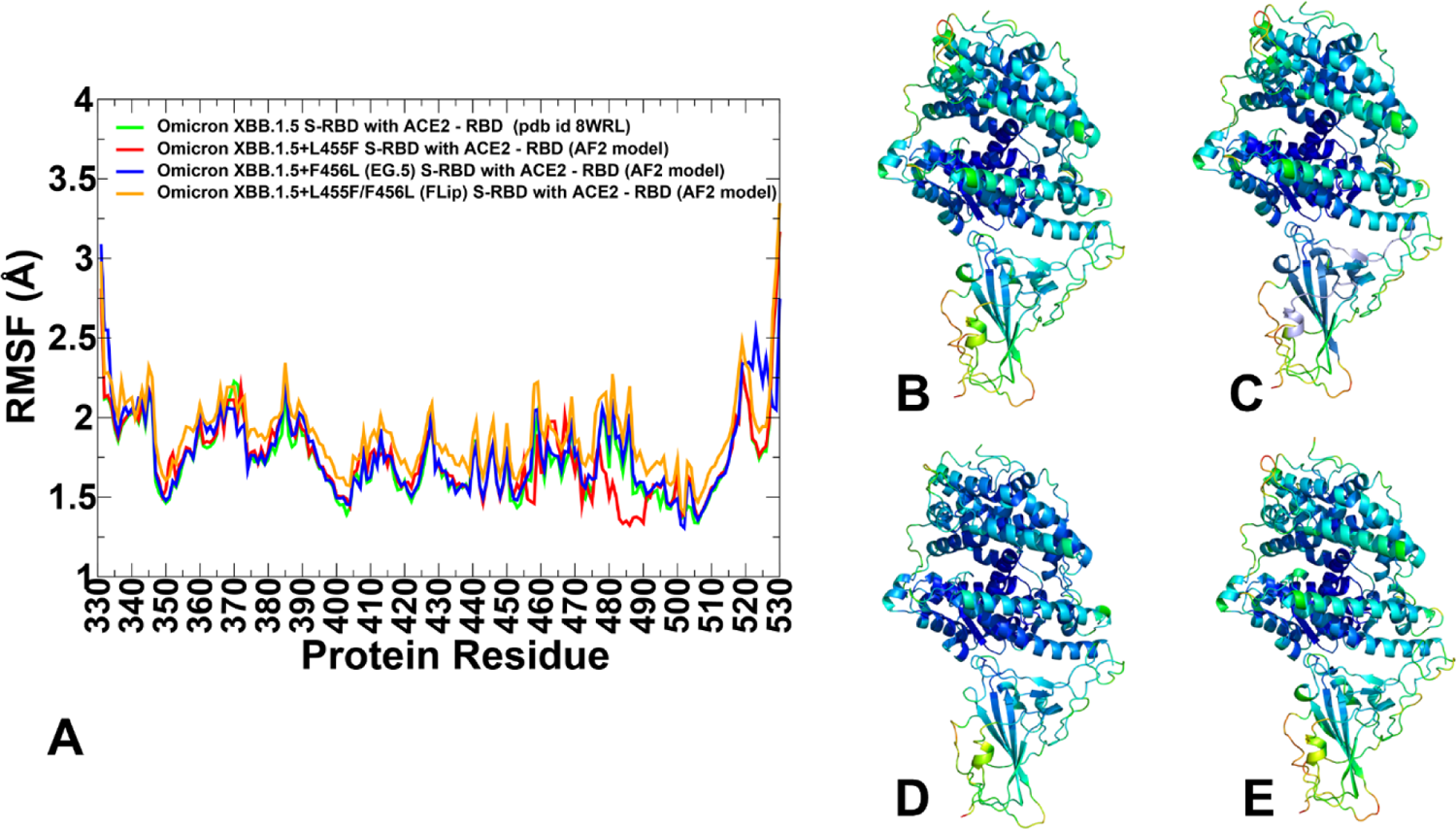
Conformational dynamics profiles obtained MD simulations of the XBB RBD-ACE2 complexes. (A) The RMSF profiles for the RBD residues obtained from MD simulations of the XBB.1.5 RBD-ACE2 complex, pdb id 8WRL (in green lines), XBB.1.5+L455F RBD-ACE2 complex, AF2 best model (in red lines), XBB.1.5+F456L RBD-ACE2 complex, AF2 best model (in blue lines), and XBB.1.5+L455F/F456L FLip RBD-ACE2 complex, AF2 best model (in orange lines). Structural mapping of the conformational dynamics profiles for the XBB.1.5 RBD-ACE2 complex (B), XBB.1.5+L455F RBD-ACE2 complex (C), XBB.1.5+F456L RBD-ACE2 complex (D) and XBB.1.5 FLip RBD-ACE2 complex (E). The profiles are mapped with the rigidity-flexibility sliding scale colored from blue (most rigid) to red (most flexible).

**Table 1.**
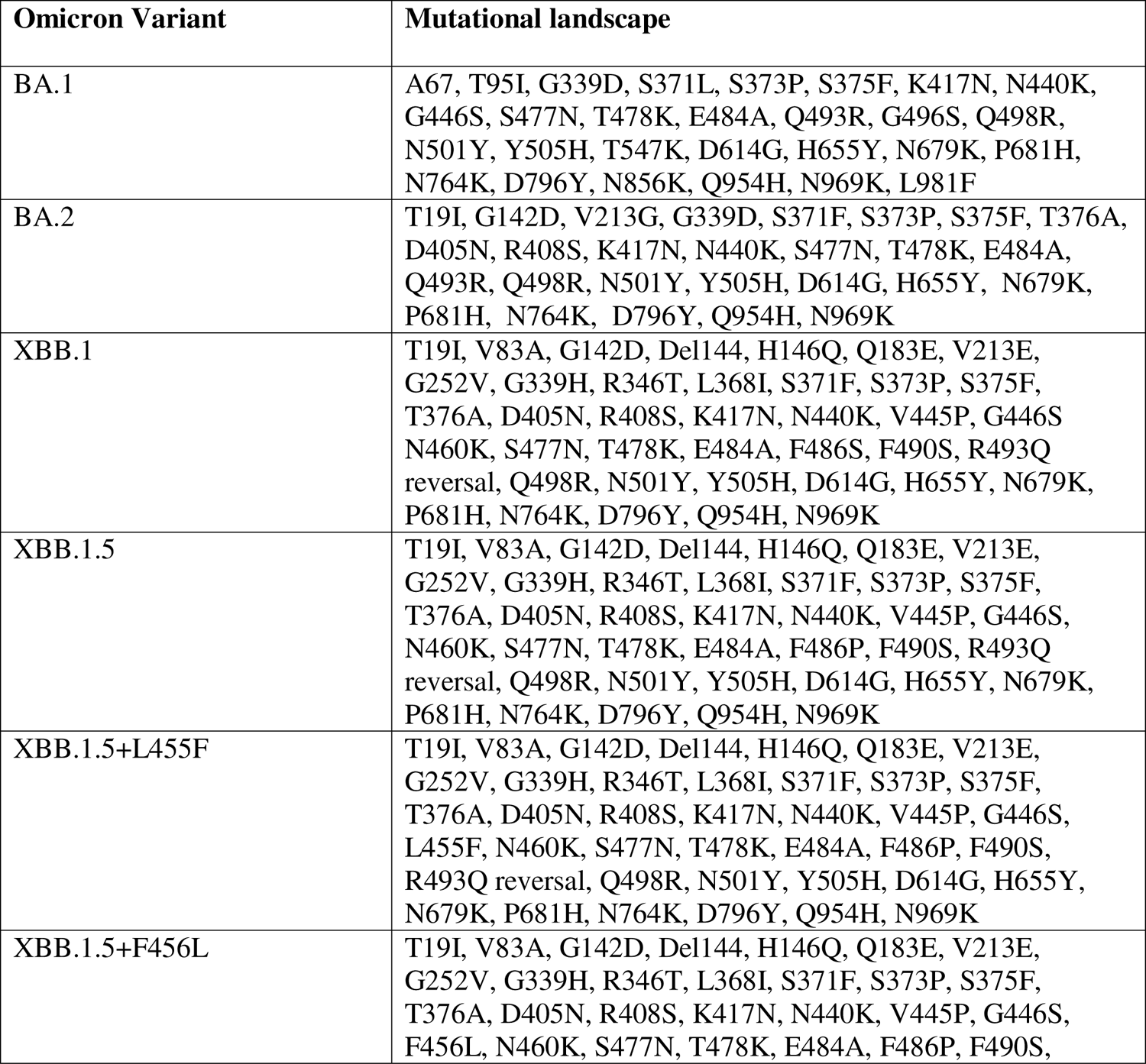
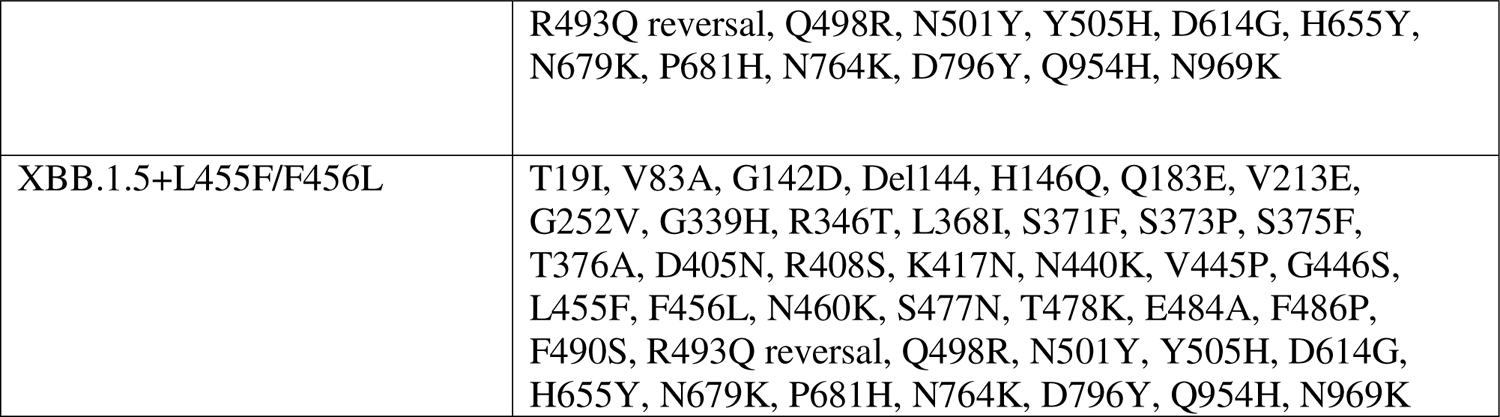
Mutational landscape of the Omicron BA.1, BA.2, XBB.1.5 variants.

| Omicron Variant | Mutational landscape |
| --- | --- |
| BA.1 | A67, T95I, G339D, S371L, S373P, S375F, K417N, N440K, G446S, S477N, T478K, E484A, Q493R, G496S, Q498R, N501Y, Y505H, T547K, D614G, H655Y, N679K, P681H, N764K, D796Y, N856K, Q954H, N969K, L981F |
| BA.2 | T19I, G142D, V213G, G339D, S371F, S373P, S375F, T376A, D405N, R408S, K417N, N440K, S477N, T478K, E484A, Q493R, Q498R, N501Y, Y505H, D614G, H655Y, N679K, P681H, N764K, D796Y, Q954H, N969K |
| XBB.1 | T19I, V83A, G142D, Del144, H146Q, Q183E, V213E, G252V, G339H, R346T, L368I, S371F, S373P, S375F, T376A, D405N, R408S, K417N, N440K, V445P, G446S, N460K, S477N, T478K, E484A, F486S, F490S, R493Q reversal, Q498R, N501Y, Y505H, D614G, H655Y, N679K, P681H, N764K, D796Y, Q954H, N969K |
| XBB.1.5 | T19I, V83A, G142D, Del144, H146Q, Q183E, V213E, G252V, G339H, R346T, L368I, S371F, S373P, S375F, T376A, D405N, R408S, K417N, N440K, V445P, G446S, N460K, S477N, T478K, E484A, F486P, F490S, R493Q reversal, Q498R, N501Y, Y505H, D614G, H655Y, N679K, P681H, N764K, D796Y, Q954H, N969K |
| XBB.1.5+L455F | T19I, V83A, G142D, Del144, H146Q, Q183E, V213E, G252V, G339H, R346T, L368I, S371F, S373P, S375F, T376A, D405N, R408S, K417N, N440K, V445P, G446S, L455F, N460K, S477N, T478K, E484A, F486P, F490S, R493Q reversal, Q498R, N501Y, Y505H, D614G, H655Y, N679K, P681H, N764K, D796Y, Q954H, N969K |
| XBB.1.5+F456L | T19I, V83A, G142D, Del144, H146Q, Q183E, V213E, G252V, G339H, R346T, L368I, S371F, S373P, S375F, T376A, D405N, R408S, K417N, N440K, V445P, G446S, F456L, N460K, S477N, T478K, E484A, F486P, F490S, |
|  | R493Q reversal, Q498R, N501Y, Y505H, D614G, H655Y, N679K, P681H, N764K, D796Y, Q954H, N969K |
| XBB.1.5+L455F/F456L | T19I, V83A, G142D, Del144, H146Q, Q183E, V213E, G252V, G339H, R346T, L368I, S371F, S373P, S375F, T376A, D405N, R408S, K417N, N440K, V445P, G446S, L455F, F456L, N460K, S477N, T478K, E484A, F486P, F490S, R493Q reversal, Q498R, N501Y, Y505H, D614G, H655Y, N679K, P681H, N764K, D796Y, Q954H, N969K |

**Table 2.**
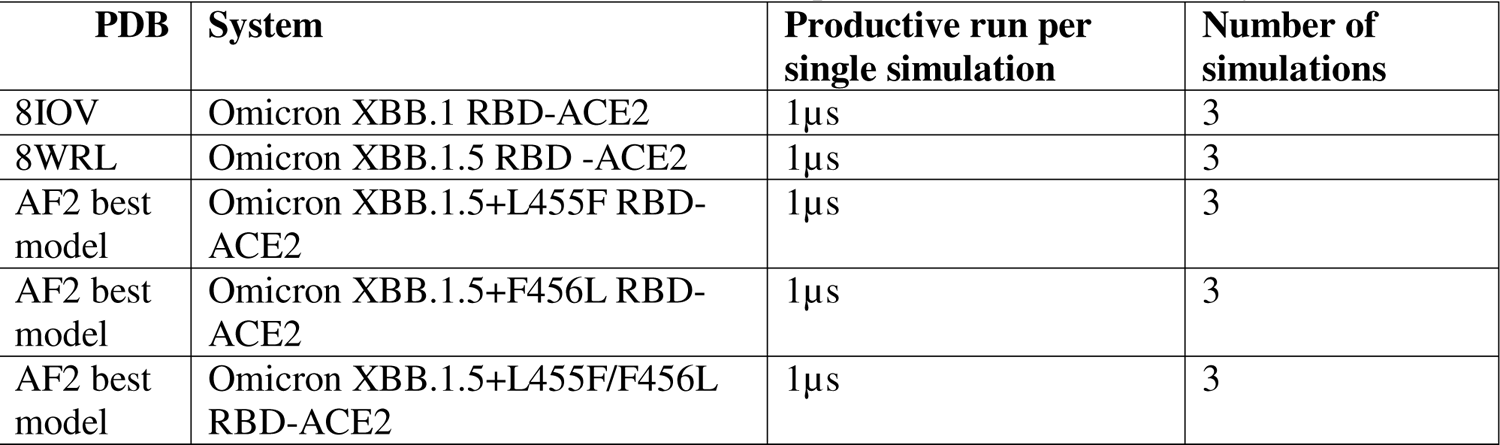
Structures of the Omicron RBD-ACE2 complexes simulated in this study.

We first performed comparative all-atom MD simulations of the XBB.1 RBD-ACE2 and XBB.1.5 RBD-ACE2 complexes. For the XBB.1.5+L455F, XBB.1.5+F456L and FLip variants, the initial structures for MD simulations corresponded to the respective best AF2 models. Conformational dynamics profiles obtained from MD simulations were similar and revealed several important trends (Figure 7). The RMSF profiles showed local minima regions corresponding to the structured five-stranded antiparallel β-sheet core region that functions as a stable core scaffold (residues 350-360, 375-380, 394-403) and the interfacial RBD positions involved in the contacts with the ACE2 receptor (Figure 7A). The RMSF profiles exhibited a similar overall shape featuring common RMSF peaks corresponding to the flexible RBD regions (residues 360-373 and residues 380-396) (Figure 7A). The stable RBD core regions (residues 390-420, 430-450) exhibited small fluctuations in all complexes suggesting the increased RBD stability for these variants which may be a relevant contributing factor to the stronger ACE2 binding experimentally observed for XBB.1.5 subvariants. In addition, the RMSF profiles are characterized by several local minima corresponding to the ACE2 interfacial sites (residues 485-505) (Figure 7A). Consistent with AF2 predictions, the dynamics profiles across all XBB variants displayed displacements in the flexible RBD regions (residues 355-375, 381-394, 444-452, 455-471, 475-487). Despite profound similarities of the RMSF profiles, we found several notable differences that are particularly important in the context of the AF2 predictions. Specifically, a single additional substitution L455F in the XBB.1.5 variant appeared to rigidify the immediate proximity of the mutational site, and even more importantly markedly reduce the flexibility of the RBM tip region (residues 475-490) (Figure 7A). This is consistent with the AF2-based ensemble predictions showing a dominant contribution of the native XBB.1.5 RBD conformation and reduction of alternative flexible substates peaks, which suggested a more rigid ensemble of the RBD conformations for this subvariant (Figure 4C, 7A). The observed congruence between the AF2 and MD generated conformational ensembles and their key features also suggested a possibility of communication between functional residues L455/F456 and dynamic changes inflicted on the RBM tip region harboring mutational sites E484A and F486P.

It is widely believed that XBB mutations in the RBM tip may have evolved to enhance the conformational heterogeneity in this region and promote stronger immune escape while only moderately sacrificing AE2 binding.^41–47^ As a result, it may be suggested that individually L455F would not significantly alter the antibody neutralization profile as compared to parental XBB.1.5. Indeed, recent functional studies showed that the extent of antibody escape for XBB.1.5+L455F was largely comparable to its parental XBB.1.5 variant.^46^ At the same time, we noticed that individual F456L mutation in the EG.5 (XBB.1.5+F456L) variant mostly conforms to the XBB.1.5 dynamics profile and restores the mobility level of RBM seen in the parental variant (Figure 7A). Interestingly, in the FLip variant, a combination L455F and F456L mutation could confer the moderately increase mobility distributed over the entire RBD and particularly for flexible regions in the vicinity of 455/456 positions as well as in the RBM tip region (Figure 7A). Importantly, these results echoed the analysis of AF2-derived conformational ensembles revealing similar dynamic signatures and suggesting that FLip mutations may increase conformational heterogeneity and mobility of the RBM loops (Figure 7).

Based on these findings, one could also suggest that the plasticity of these regions may contribute to the experimentally observed synchronous effect of L455F and F456L on increasing antibody escape as compared to the XBB.1.5 as well as XBB.1.5+L455F and XBB.1.5+F456L variants.^46^ Despite the increased mobility, the RBM tip maintains a stable folded “hook” conformation that is similar to the cryo-EM conformations of XBB.1 and XBB.1.5 RBD. Our previous studies showed that well-ordered and stable “hook-like” conformation of the RBM tip is maintained in BA.2 and XBB.1.5 variants due to hydrophobic interactions provided by F486 (BA.2) and F486P (XBB.1.5).^76,77^

### Ensemble-Based Mutational Scanning of the RBD Binding Residues and Epistatic Relationships Between Binding Energy Hotspots in the XBB Lineages

Conformational ensembles obtained from MD simulations were used to perform a systematic ensemble-based mutational scanning of the RBD residues in the XBB RBD-ACE2 complexes (Figure 8). The mutational scanning heatmaps highlighted the spectrum of binding free energy changes in the RBD interface residues that maintain stable contacts with the ACE2 receptor during simulations. First, we noticed a consistent presence of shared binding energy hotspots across the XBB RBD-ACE2 complexes that correspond to the hydrophobic centers Y453, L455, F456, Y489 and Y501 (Figure 8). The heatmaps showed that Y489 and Y501 hotspots are particularly sensitive to modifications across all XBB variants as most of the substitutions in these positions induce significant destabilizing effects with ΔΔG > 2.0 kcal/mol (Figure 8). Notably, however, mutational maps showed that the increased hydrophobicity in these positions, for example Y453F/W, L455F/W may result in neutral or modestly favorable binding changes^125^ (Figure 8).

**Figure 8.**
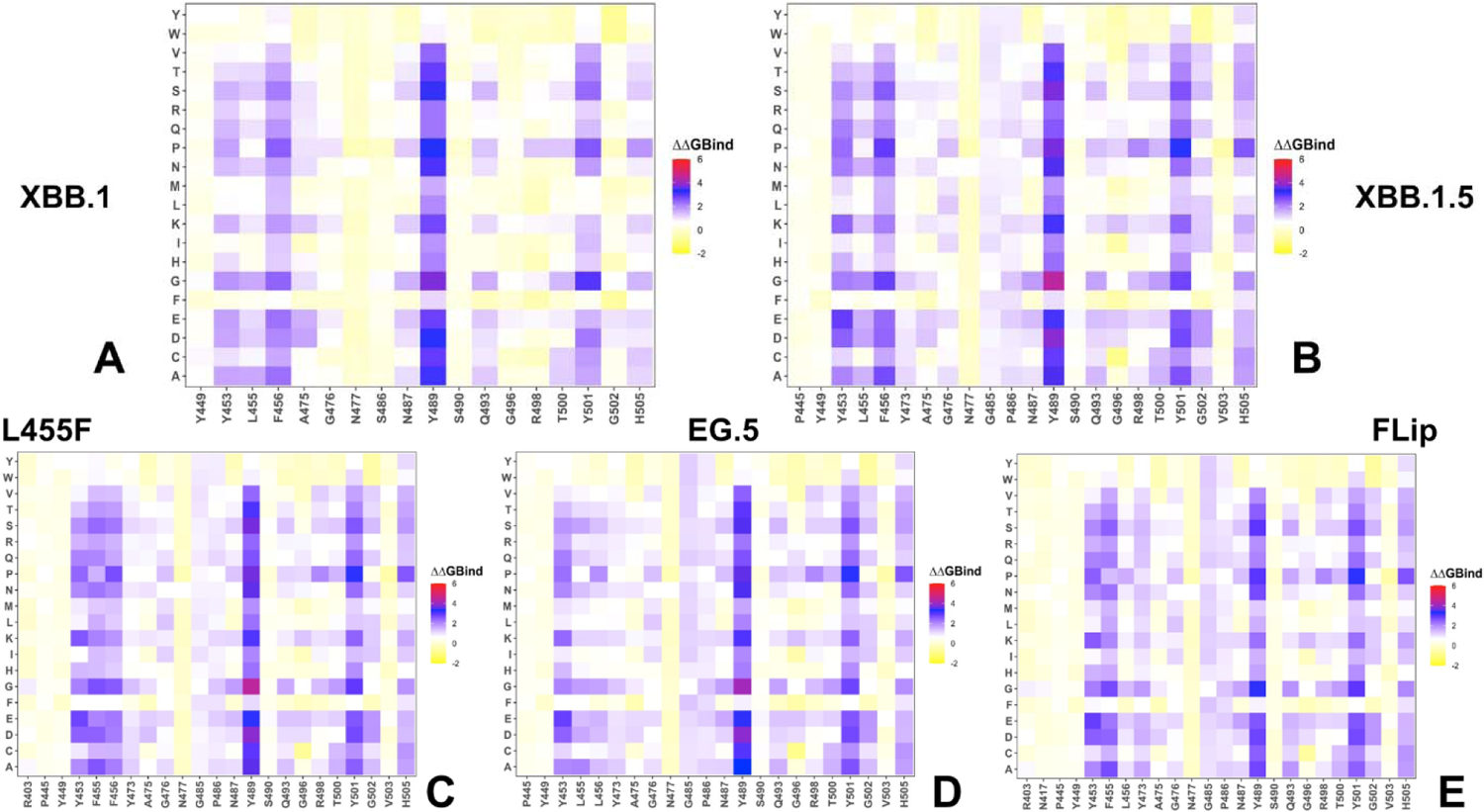
Mutational profiling of the RBD intermolecular interfaces in the Omicron RBD-ACE2 complexes. The mutational scanning heatmaps are shown for the interfacial RBD residues in the XBB.1 RBD-ACE2 complex (A), XBB.1.5 RBD-ACE2 complex (B), XBB.1.5+L455F RBD-ACE2 complex (C), XBB.1.5+F456L RBD-ACE2 complex (D) and XBB.1.5+L455F/F456L FLip RBD-ACE2 complex (E).The standard errors of the mean for binding free energy changes and are within ∼ 0.11-0.15 kcal/mol using averages based on a total of 1,000 samples obtained from the three MD trajectories for each complex.

Interestingly, the three adjacent RBD hotspots Y453, L455 and F456 located within the central strands comprising the ACE2 interface showed appreciable epistatic shifts between BA.2 and BQ.1.1 or XBB.1.5, including prominent epistatic changes in the effects of mutations Y453W, L455W/F and F456L depending on the genetic background.^62^ Previous DMS experiments also demonstrated that Y453, L455, F456, F486, and Y489 play a fundamentally critical role in determining both structural stability of the RBD fold and binding with ACE2.^54–56^ These conserved residues form consistent and similar interactions in all Omicron variants, and mutations in some of these positions are typically conservative allowing for preservation of their key functional role as determinants of the RBD stability. ^125^ The mutational heatmap of the XBB.1.5 Flip RBD residues highlighted the increased stabilization role of Y453, F455 and L456 “trio” of residues (Figure 8E) as L455F/F456L mutations appeared to enhance the contribution of these three residues for binding with ACE2. Previous DMS experiments quantified the effects of F486 mutations (F486V/I/S/L/A/P) showing that even though substitutions of F486 can reduce binding F486P imposes the lowest cost in RBD affinity loss and has the largest increase in RBD expression.^54–56,60^

Mutational heatmaps showed that mutations in the S486 position of XBB.1 (Figure 8A) are more forgiving as compared to scanning of P486 in the XBB.1.5 variant (Figure 8B) which is consistent with the more favorable ACE2 binding upon S486P modifications. The heatmaps of the XBB.1.5+F456L (Figure 8D) and FLip variants (Figure 8E) demonstrated that mutations of P486 to positions other than reversed P486F are typically only moderately destabilizing and are unlikely to impose significant binding loss. Consistent with that, it was found that this position exemplified by F486V (BA.4/BA.5), F486I, F486S (XBB.1), and F486P mutational changes (XBB.1.5, EG.5, EG.5.1, FLip) represents a convergent evolutionary hotspot and is one of the major hotspots for escaping neutralization by antibodies.^39^ Another critical site of convergent evolution is reversed R493Q mutation and mutational scanning results showed that modifications at Q493 positions are generally destabilizing. This is consistent with the notion that R493Q may compensate for partial binding loss incurred by F486P mutation. These findings are also in agreement with previous studies showing that R493Q reversal may induce the increased affinity of the RBD with ACE2 receptor.^127^

To better understand the functional effects of the XBB.1.5 mutations and evaluate potential epistatic relationships between the RBD hotspots, we computed binding free energies induced by the XBB.1.5 RBD mutations (Figure 9). Notably, some of the XBB.1.5 mutational sites are not in direct contact with ACE2, and the computed changes reflect cumulative effect of both structural stability and binding affinity effects. In addition, we also included Y453W, L455F and F456L mutations which allowed for head-to-head comparisons between different XBB lineages and quantified the epistatic couplings between mutational sites (Figure 9). First, we consistently observed that most of the XBB.1.5 mutations (including G339H R346T, L368I, S371F, S373P, S375F, T376A, D405N, R408S, K417N, N440K, V445H, G446S, N460K, S477N, T478K and E484A) cause only very moderate and stabilizing changes (Figure 9). Although the cumulative effect of these stabilizing changes may add up and provide a meaningful contribution to ACE2 binding, these mutations play a marginal role in determining the ACE2 binding. Instead, these mutations are exploited by the virus to enhance and manipulate the immune evasion profile and enable resistance to neutralization against distinct classes of antibodies.

**Figure 9.**
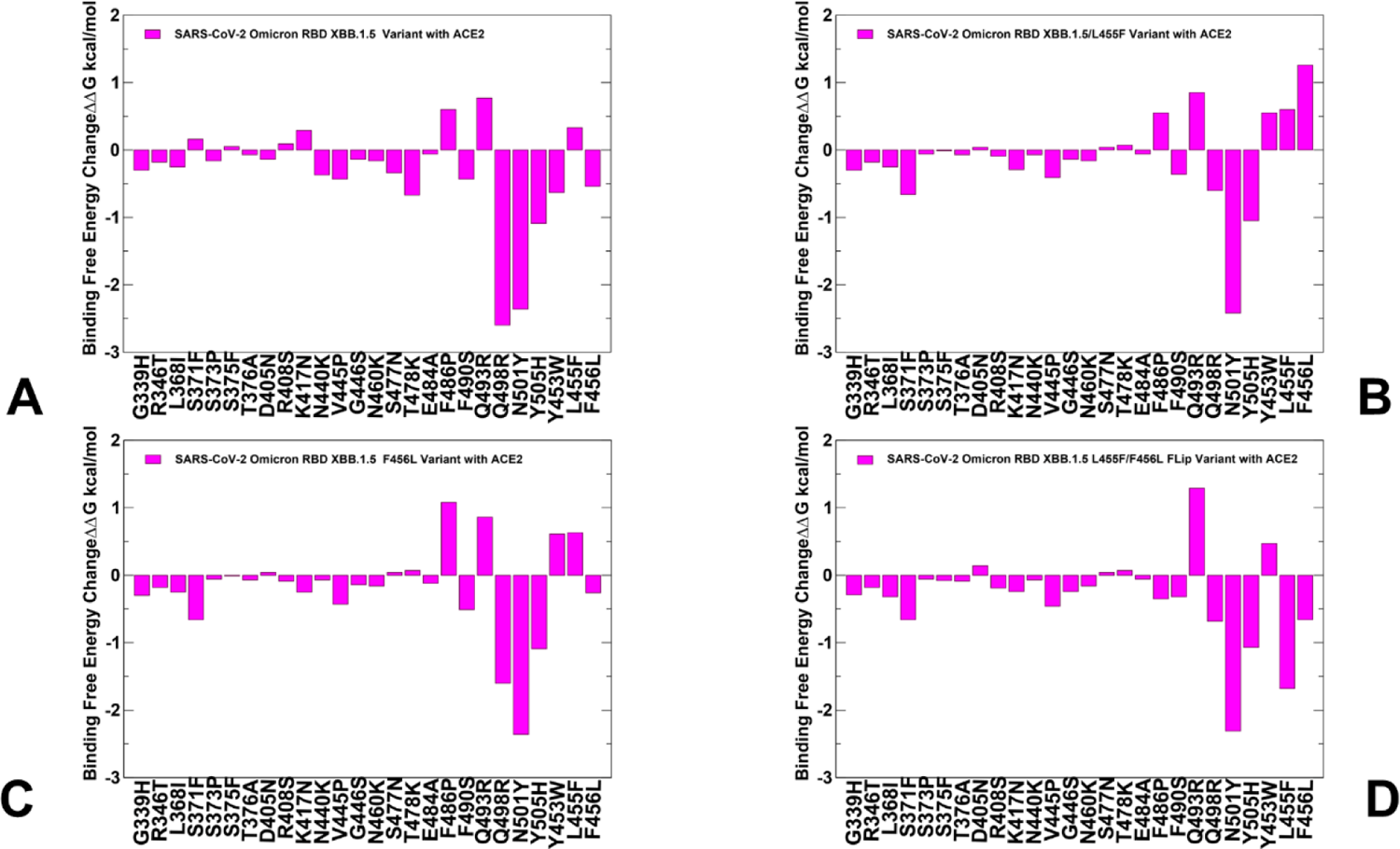
The ensemble-based binding free energy changes obtained from MD simulations for the XBB.1.5 mutations induced in different genetic backgrounds of the XBB.1.5 variant (A), XBB.1.5+L455F variant (B). XBB.1.5+F456L variant (C) and XBB.1.5+FLip variant (D). The binding free energies are shown in magenta-colored filled bars. The positive binding free energy values ΔΔG correspond to destabilizing changes and negative binding free energy changes are associated with stabilizing changes.

Another common feature is that mutations Q498R, N501Y and Y505H which are present in other Omicron variants make the largest contributions to the binding affinity and correspond to the major binding affinity hotspots. The intriguing aspect of this comparative analysis is the observed differences in the effects of XBB.1.5 mutational sites including F486P, F490S, Q493R as well as L455F and F456L computed in the XBB.1.5 background (Figure 9). First, we found that the destabilizing effect of F486P is small, particularly in the XBB.1.5 and XBB.1.5+L455F variants (Figure 9A,B) and yet varies between XBB lineages to become even marginally stabilizing in the FLip variant (Figure 9D). This may result from the increased plasticity of the RBM tip that allows for the greater number of favorable P486 interactions with ACE2 and potentially entropy-mediated improvement in the binding. Strikingly, we also observed changes in the effects of L455F and F456L mutations. The binding effect of L455F mutation in the XBB.1.5, XBB.1.5+L455F and XBB.1.5 F456L backgrounds is moderately destabilizing (Figure 9A-C) but becomes appreciably favorable when introduced in combination with F456L in the FLip variant. In addition, we also found that Y453W and F456L are marginally stabilizing in the XBB.1.5 (Figure 9A) which is fully consistent with the DMS experiments showing that these mutations may enhance ACE2-binding affinity by 7.1- and 1.9-fold in XBB.1.5 background.^62^

The central result of this analysis is the markedly increased stabilization effect of Q493 in the FLip variant and corresponding synchronization in favorable contributions of L455F and F456L mutations (Figure 9D). These observations suggested a coordinated epistatic effect between Q493 and L455F/F456L doubled mutation that is observed only in the FLip variant. In this context, our results corroborate the functional experiments^46^ and DMS studies that measured ACE2-binding affinities of the Y453W, L455F, and F456L mutations in combination with Q493R or R493Q in the XBB.1.5 and BA.2 backgrounds.^62^ These experiments found a striking epistatic shift where L455F caused strong affinity reduction on XBB.1.5, but significantly improved the ACE2-binding affinity of XBB.1.5+F456L.^46^ Our computations reproduced this effect, particularly showing that the L455F contribution to binding is completely reversed and becomes highly favorable when paired with F456L mutation (Figure 9D). Our analysis also corroborated with the proposed role of Q493 as mediator of epistatic interactions as the increased favorable contribution of Q493 in the FLip variant may be linked with the concreted effect of L455F/F456L pair.

To examine global predictions of in silico mutational scanning, we compared the results of the DMS experiments for the BA.2 and XBB.1.5 RBD residues^54–56,62^ with the computed mutational changes in the BA.2 RBD-ACE2 ((Supporting Information, Figure S5A) and XBB.1.5 RBD-hACE2 complexes (Supporting Information, Figure S5B). A statistically significant correlation between the DMS experiments and mutational scanning data was observed, particularly for BA.2 RBD. also highlighting a significant dispersion of the distributions. It is worth noting that the computed free energy changes reflected mutation-induced effects on both residue stability and binding interactions. It could be noticed that the computational predictions of destabilizing changes were often larger than the experimentally observed values. Nonetheless, the scatter plots showed a fairly appreciable correspondence between the predicted and experimental free energy differences, particularly for large destabilizing changes with ΔΔG > 2.0 kcal/mol ((Supporting Information, Figure S5A-C). This ensures robust identification of major binding affinity hotspots where mutations cause pronounced energetic changes. We also compared the DMS free energies for the XBB.1.5 RBD mutational sites only with the computationally predicted binding changes with ACE2 (Supporting Information, Figure S5D), showing an excellent correspondence between the computational and experimental data.

Recent evolutionary studies suggested that during evolution there is the cumulative effect of many small-effect epistatic modifications, and in the background of gradual epistatic drifts, a few mutations may occasionally undergo substantial changes in their effects.^128,129^ While extensive epistatic shift was discovered between Q498R and N501Y mutation combination^57,58^, the epistatic interactions between Q493, L455F and F456L may be more gradual. These gradual epistatic changes may emerge to mediate strong ACE2 binding by amplifying contributions of a small number of binding hotspots, while deploying mutations in other positions to combat antibody binding. Within XBB, FLip lineages continue to grow and A475V continues to be beneficial when combined with L455F/F456L FLip.^3^ Interestingly, with XBB acquiring multiple mutations since its emergence, we increasingly observe lineages with pair or triplets of RBD mutations, for example pairing F456L and K478R that emerged as another beneficial combination that can confer growth advantage for the virus.^39^

A growth advantage of the combination F456L + K478R over either mutation alone (∼35% per week, doubling every week), suggests that more lineages with this combination could emerge in the immediate future.^39^ In this context, MD simulations and mutational scanning results indicated that L455F and F456L mutations may differentially affect the heterogeneity of the RBM tip that harbors K478 position. According to our results, the FLip variant may increase the flexibility of the RBM loop and mediate further heterogeneity upon K478R mutation which may be potentially exploited by the virus to boost immune evasion potential.

### MM-GBSA Analysis of the Binding Affinities for the XBB RBD-ACE2 Complexes: Energetics of Binding Energy Hotspots and FLip Variant-Induced Enhanced ACE2 Binding

Using the conformational equilibrium ensembles obtained from MD simulations, we computed the binding free energies for the XBB RBD-ACE2 complexes using the MM-GBSA method^109,110^ A common strategy to reduce noise and cancel errors in MM-GBSA computations is to run MD simulations on the complex only, with snapshots taken from a single trajectory to calculate each free energy component. The results of MM-GBSA computations showed excellent agreement with the experimental SPR-measured binding affinities for the XBB RBD-ACE2 complexes (Table 3, Figures 10,11A). Consistent with the experiments, we found that FLip mutations not only restore the binding strength of the parental XBB.1.5 variant but could also lead to moderate improvement in binding affinity (Table 3).

**Figure 10.**
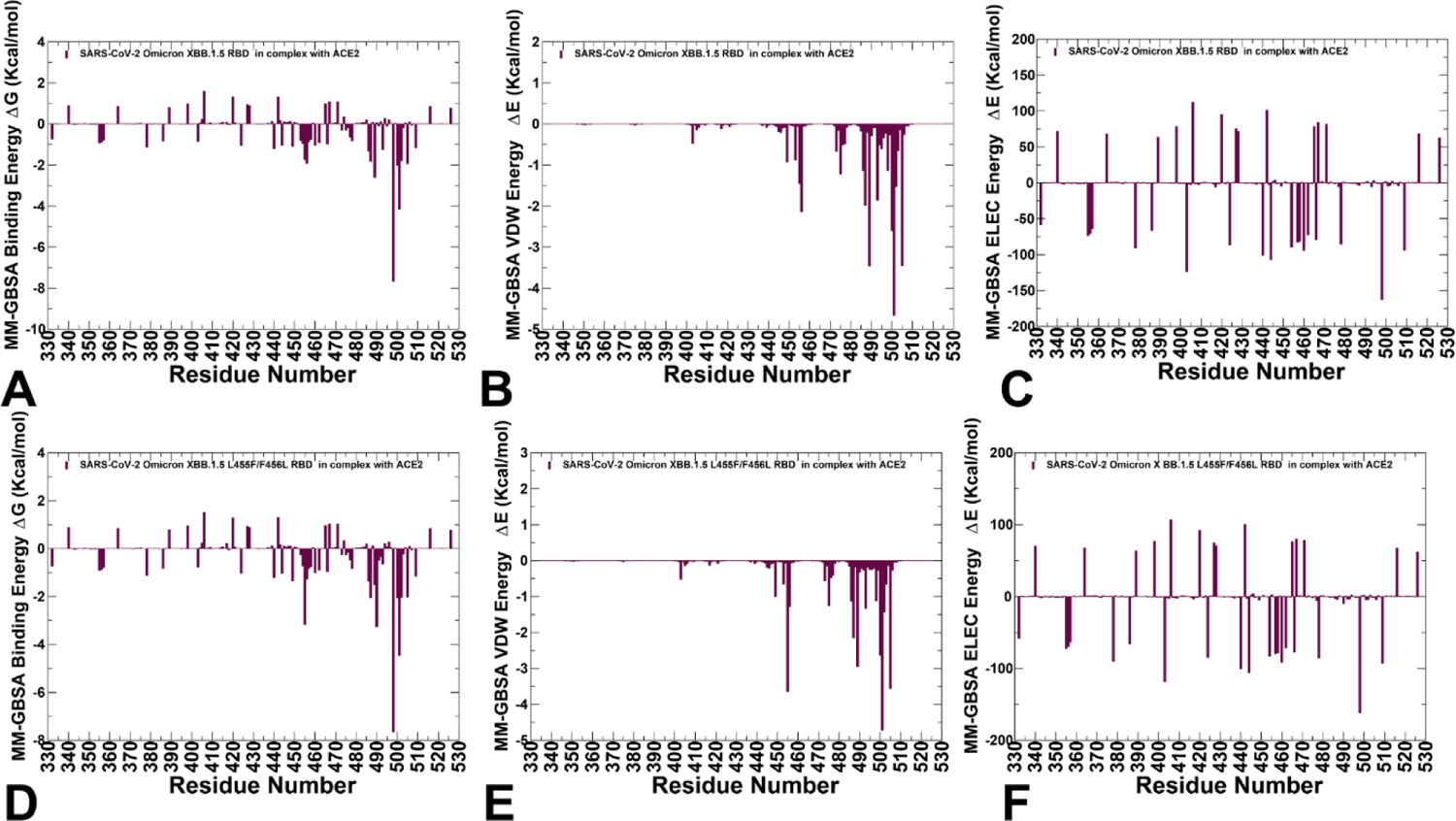
The residue-based decomposition of the MM-GBSA energies. (A) The residue-based decomposition of the total MM-GBSA binding energy ΔG contribution for the XBB.1.5 RBD-ACE2 complex. (B) The van der Waals contribution of the total binding energy for the XBB.1.5 RBD-ACE2 complex. (C) The electrostatic contribution of the total binding energy for the XBB.1.5 RBD-ACE2 complex. The residue-based decomposition of the total MM-GBSA binding energy ΔG contribution for the XBB.1.5 FLip RBD-ACE2 complex (D), the van der Waals contribution of the XBB1.5 FLip total binding energy (E) and the electrostatic contribution of the XBB.1.5 FLip total binding energy (F). Th MM-GBSA contributions are evaluated using 1,000 samples from the equilibrium MD simulations of respective RBD-ACE2 complexes. It is assumed that the entropy contributions for binding are similar and are not considered in the analysis. The statistical errors was estimated on the basis of the deviation between block average and are within 0.25-0.95 kcal/mol.

**Figure 11.**
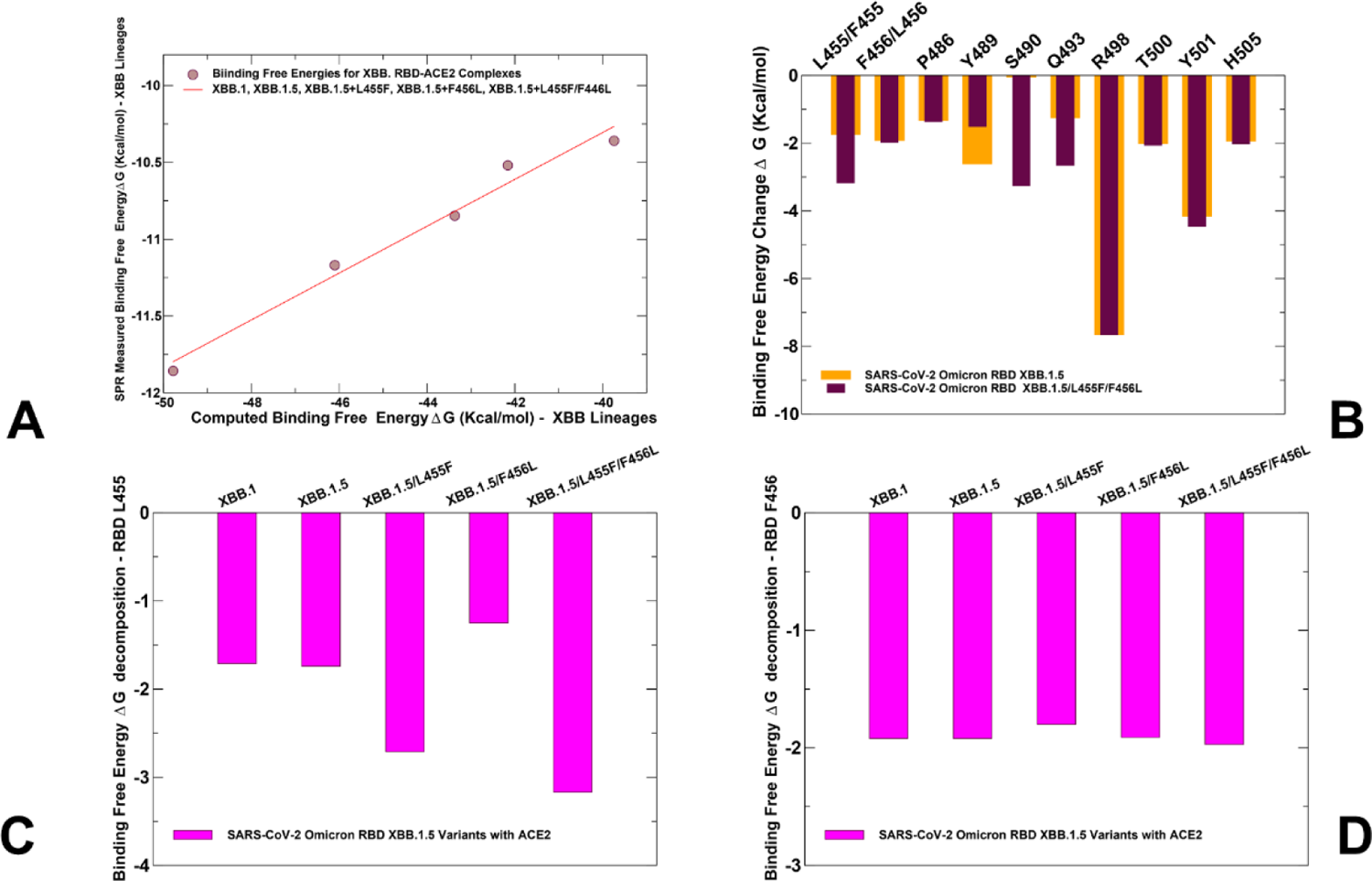
The MM-GBSA analysis of binding affinity changes in the XBB RBD-ACE2 complexes. (A) The scatter correlation plot of the experimental binding affinities and MM-GBSA total binding energies for the XBB.1, XBB.1.5, XBB.1.5+L455F, XBB.1.5+F456L and XBB.1.5 FLip RBD-ACE2 complexes. (B) A comparison of the MM-GBSA binding free energy contributions between the XBB.1.5 and XBB.1.5 FLip RBD-ACE2 complexes. The orange bars (XBB.1.5) and maroon-colored bars (XBB.1.5 FLip) represent MM-GBSA contributions for residues L455/F455, F456/L456, P486, Y489, S490, Q493, R498, T500, Y501, and H505. (C,D) A comparison of the MM-GBSA binding energy contributions for L455 (F455) residue and F456 (L456) residue in the XBB.1, XBB.1.5, XBB.1.5+L455F, XBB.1.5+F456L and XBB.1.5 FLip complexes. The respective contributions (magenta-colored filled bars) are obtained from MM-GBSA decomposition analysis.

**Table 3.** MM-GBSA Binding Energies for the XBB RBD-ACE2 Complexes.

| System | $E_{vdW}$ | $E_{elec}$ | $E_{GB}$ | $E_{SA}$ | $\Delta G_{bind}$<br>(kcal/mol) | Exp.binding |
| --- | --- | --- | --- | --- | --- | --- |
| XBB.1 RBD-ACE2 | -70.72 | -1241.68 | 1280.04 | -9.8 | -42.16 | 19nM/ |
| XBB.1.5 RBD-ACE2 | -74.41 | -1236.49 | 1274.50 | -9.7 | -46.1 | 6.4 nM |
| XBB.1.5+L455F RBD-ACE2 | -73.33 | -1225.64 | 1268.80 | -9.56 | -39.74 | 25 nM |
| XBB.1.5+F456L RBD-ACE2 | -70.61 | -1233.02 | 1269.03 | -8.88 | -43.37 | 11nM |
| XBB.1.5_L455F/F456L RBD-ACE2 | -74.01 | -1250.34 | 1284.35 | -9.77 | -49.78 | 2nM |

Interestingly, the decomposition of binding free energy terms showed very similar contributions of the van der Waals interactions in the XBB.1. 5 and XBB.1.5 FLip RBD-ACE2 complexes, while the main determinant of the improved binding is the more favorable electrostatic interactions in the FLip complex (Table 3). At the same time, we observed that for XBB.1.5+L455F variant the electrostatic interactions are significantly weakened as compared to the parental XBB.1.5 while the van der Waals contacts remain as favorable as for XBB.1.5 (Table 3). In some contrast, F456L mutation could moderately weaken both the nonpolar and electrostatic interactions, resulting in the partial loss of binding affinity compared to XBB.1.5 variant (Table 3).

Our structural analysis suggested that in the XBB.1.5 RBD-ACE2 complex, Q493 makes fairly weak contacts with H34 and K31 ACE2 binding residues as the corresponding average distances of Q493-H34 are ∼ 4.5 Å and Q493-K31 are ∼ 4.3 Å ( Figure 1E). On the other hand, Q493 makes better interactions in the FLip variant where the average distances of Q493-H34 are ∼ 3.6 Å and Q493-K31 are ∼ 4.1 Å respectively, while H34 side chain can also establish favorable contacts with S494 within ∼3.8 Å (Figure 1H). These structural predictions of the ensemble-averaged XBB RBD-ACE2 complexes are consistent with the latest experimental evidence that rearrangement in the binding interface of the FLip RBD-ACE2 complex allow for more favorable interactions of H34 with both Q493 and S494 RBD residues.^46^

The residue-based decomposition of the MM-GBSA energies is particularly revealing when comparing the MM-GBSA breakdown for XBB.1.5 RBD-ACE2 complex (Figure 10A-C) and the XBB.1.5+FLip variant complex (Figure 10D-F). The major RBD contributors of binding affinity in the XBB RBD-ACE2 complexes are residues L455, F456, P486, N487, Y489, Q493, R498, T500, Y501 and G502. Importantly, the strongest binding hotspots correspond to universal Omicron hotspots R498 and Y501 that are also connected via strong epistatic interactions^57,58^ and collectively drive binding affinity of the RBD-ACE2 complexes (Figure 10). In addition, the favorable binding contributions are also provided by sites K378, R403, K424, K440, K444, Y453, K460, N477, K478 (Figure 10). Similar trends were also observed in residue-based decomposition of MM-GBSA energies for XBB.1.5+L455F and XBB.1.5+F456L variants (Supporting Information, Figure S6).

The individually moderate but cumulatively significant contribution of these residues is determined primarily by strong electrostatic interactions mediated by lysine residues, which is the result of Omicron evolution leading to significant accumulation of positively charged substitutions interacting with the negatively charged ACE2 binding interface residues. Recent studies mapped the electrostatic potential surface of S protein and major variants to show accumulated positive charges at the ACE2-binding interface, revealing the critical role of complementary electrostatic interactions driving the enhanced affinity the Omicron S-ACE2 complexes.^130,131^ Extensive simulations investigated the electrostatic features of the RBD for the several S variants and binding affinities at several pH regimes revealing that the virus evolution may primarily exploit the electrostatic forces to make RBD more positively charged and improve the RBD-ACE2 binding affinity.^132^ It is possible that evolution of the electrostatic RBD surface in XBB Omicron variants may be exploited to enable both the incremental improvements in the ACE2 binding and resistance to antibody interactions. Of particular interest are the binding affinity contributions of L455, F456 in XBB.1.5 and their respective mutants F455/L456 in the XBB.1.5 FLip variant (Figure 11). The residue decomposition of the MM-GBSA energies showed that total contribution of L455F ( ΔG = −3.17 kcal/mol) and F456L (ΔG =-1.97 kcal/mol) in the FLip variant are more favorable than the respective contributions of L455 (ΔG =-1.74 kcal/mol) and F456 (ΔG =-1.92 kcal/ mol) in the parental XBB.1.5 (Figure 11A,B). Our results highlighted a significant role of L455F substitution that is amplified only when combined with F456L mutation which may function as a trigger of epistatic interactions in the FLip variant. This finding supports the proposed mechanism in which F456L is considered a prerequisite for the fitness of the FLip variant and mediator of the synergistic enhancement in the ACE2 binding. Another important residue mediating the improved binding for the FLip variant is Q493 that contributes ΔG =-2.65 kcal/mol, whereas in the XBB.1.5 variant the respective Q493 contribution ΔG =-1.25 kcal/mol is smaller but still favorable for ACE2 binding (Figure 11B).

Notably, the MM-GBSA decomposition analysis revealed that the favorable contributions of F486, R498, T500 and Y501 hotspots in the XBB.1.5 and FLip complexes are similar (Figure 11B). According to our results, the binding affinity differences between these XBB variants may be determined by L455F, F490S and Q493 positions (Figure 11B). A more detailed comparison of binding contributions mediated by van der Waals interactions in L455 position showed that this component becomes highly favorable in the L455F and L455F/F456L variants (Figure 11C). A close inspection of the binding energies mediated by interactions in the F456 position indicated favorable but similar contributions of F456 and F456L (Figure 11D), suggesting that F456L may function as facilitator of synergistic effects in the FLip variant rather than a single determining factor.

Together, these findings provided a useful insight into the mechanism underlying stronger binding of the XBB.1.5 FLip variant relative to XBB.1 and XBB.1.5 suggesting that differences in the binding affinity are driven by cumulative contributions and balance of electrostatic and nonpolar interactions for several important RBD hotspots including L455F/F456L, F490S, and Q493. At the same time, the universally important for binding Y489, R498 and Y501 sites maintain their strong favorable interactions in both complexes.

### Network-Based Community Analysis of Epistatic Couplings in the RBD-ACE2 Complexes

To characterize and rationalize the experimentally observed epistatic effects of the Omicron mutations we explored previously introduced a simple clique-based network model for describing the non-additive effects of the RBD residues. Using equilibrium ensembles and dynamic network modeling of the original RBD-ACE2 complexes we used mutational scanning to perturb modular network organization represented by a chain of inter-connected sable 3-cliques. Specifically, we calculated the probability by which mutational sites would belong to the same interfacial 3-clique. For this, we generated an ensemble of 1,000 protein conformations from MD simulations of the studied RBD-ACE2 complexes. To systematically estimate the non-additive effects of XBB.1.5 mutations and L455F/F456L double-site mutations, we constructed dynamic network structures for each mutant and determined the topological quantities of these networks. By using mutational changes of the Omicron positions over the course of the MD simulation trajectory for the RBD-ACE2 complexes we attribute RBD and ACE2 interfacial sites that belong to the same 3-clique to have local non-additive effects, while the effects of specific mutations on changes affected the entire distribution and the total number of 3-cliques at the RBD-ACE2 interface would be attributed to long-range epistatic relationships. If the mutational sites are arranged in a 3-clique structure, all three sites are connected to each other. As a result, when one site is mutated, it will have a greater effect on the stability of the complex because the other two sites will also be affected. Therefore, the presence of a stable 3-clique structure can be used as a first predictor of potential local non-additive effects.

We computed the distribution of stable in MD simulations 3-cliques in the XBB.1.5 RBD-ACE2 complex (Supporting Information, Figure S7) showing that R498 and Y501 sites can promote a larger number of stable 3-cliques at the central interfacial patch. In addition, Q493 participates in the following stable cliques: Q493-H34-Y453, Q493-K31-Y489, H34-K31-Q493, Q493-K31-F456, L455-H34-Q493 and Q35-Q493-H34. In addition, we detected D30-L455-F456, L455-K31-F456, Y453-T27-Y489, F456-T27-Y489 and D30-K31-F456 cliques that all share F456 site and link together F456 with L455 and D30, K31 and T27 of ACE2 (Supporting Information, Figure 7). This network-based community analysis of the XBB.1.5 RBD-ACE2 complex revealed that R498, Y501, Q493 and F456 positions mediate the vast majority of 3-cliques at the binding interface and therefore can be responsible for modulating non-additive epistatic relationships between RBD residues. The results are consistent with the DMS studies which discovered extensive epistasis between R498 and Y501 combination, while aromatic substitutions are not tolerated simultaneously at these positions.^57,58,62^ The network analysis also revealed strong and sustainable couplings in 3-cliques between Q493, Y453, L455 and F456 residues that are dominated by mediators Q493 and F456 positions (Supporting Information, Figure S7). According to the community analysis, the non-additive effects may be accentuated ensured by a chain of linked 3-cliques containing Q493 and F456 residues in which each pair of nodes/residues is connected by an edge, indicating a strong mutual interaction among amino-acids on these nodes. This also agrees with the latest DMS analysis of XBB variants showing the ongoing epistatic drift and epistatic interaction between R493Q reversion and mutations at RBD 453, 455, and 456 positions.^62^ These observations showed that all cliques at the middle portion of the interface are mediated through couplings between Q493, L455 and F456. Interestingly, the largest number of cliques include Q493 and F456 suggesting that these residues may be key mediators of non-additive couplings. The network community analysis showed that the topology of the stable interfacial cliques is preserved between XBB.1.5 and XBB.1.5 FLip complexes, but the stability of these cliques and their simulation life time can change. To quantify the degree of epistasis, we also calculated the ratio of *P_ab_* after double mutations to *P_ab_* after single mutations. If the probability of two sites belonging to the same 3-clique during simulation increases after double mutations, it would indicate that there is an epistatic effect between the two sites. We found that the simulation life time of stable 3-cliques that involve combined mutations R493Q, L455F and F456L, particularly for cliques Q493-K31-F456L, L455F-H34-Q493, D30-L455F-F456L, and L455F-K31-F456L, can increase from ∼65%-70 % for XBB.1.5 to 85%-90% for these cliques with mutated L455 and F456 in the XBB.1.5 FLip variant complex. Hence, the non-additive interactions within cliques that are mediated by R493Q and F456L may be strengthened, leading to the enhanced stability of the binding interface in the FLip RBD-ACE2 complex. Hence, a clique-based network model can identify highly correlated and potentially non-additive RBD sites and distinguish them from other mutational sites that are less likely to experience epistatic shifts.

### Mutational Profiling of Spike Protein Binding with Monoclonal RBD Antibodies: Revealing Central Role of F456L and F486P Mutations in Immune Evasion

We performed structure-based mutational analysis of the S protein binding with different classes of RBD-targeted antibodies, focusing specifically on the role of XBB.1.5 mutations as well as L455F, F456L and FLip mutations in mediating resistance to class 1 antibodies. The latest functional studies showed that EG.5 and EG.5.1 variants displayed resistance against S2K146 antibody and a more significant neutralization resistance to Omi-3, Omi-18, Omi-42 and BD-515 class 1 monoclonal antibodies.^42^ For comparison with the experimental data, we specifically examined S2K146 and Omi-3 antibodies that were experimentally tested for evasion properties against XBB.1.5+F456L (EG.5), EG.5.1, and XBC.1.6 variants.^42,43^ The structures of RBD-targeted antibodies used in this analysis included S-RBD complex with S2K146 (pdb id 7TAS)^133^ and S-RBD Omicron complex with Omi-3 (pdb id 7ZF3).^134^ Mutational profiling analysis of the RBD-antibody complexes for XBB.1.5, XBB.1.5+L455F, XBB.1.5+F456L and XBB.1.5 FLip variants (Figure 12) allowed for direct comparison with the reported fold changes in IC_50_ of antibody binding for these variants in the background of XBB.1.5 and enabled to clarify role of FLip mutations in mediating antibody resistance.

**Figure 12.**
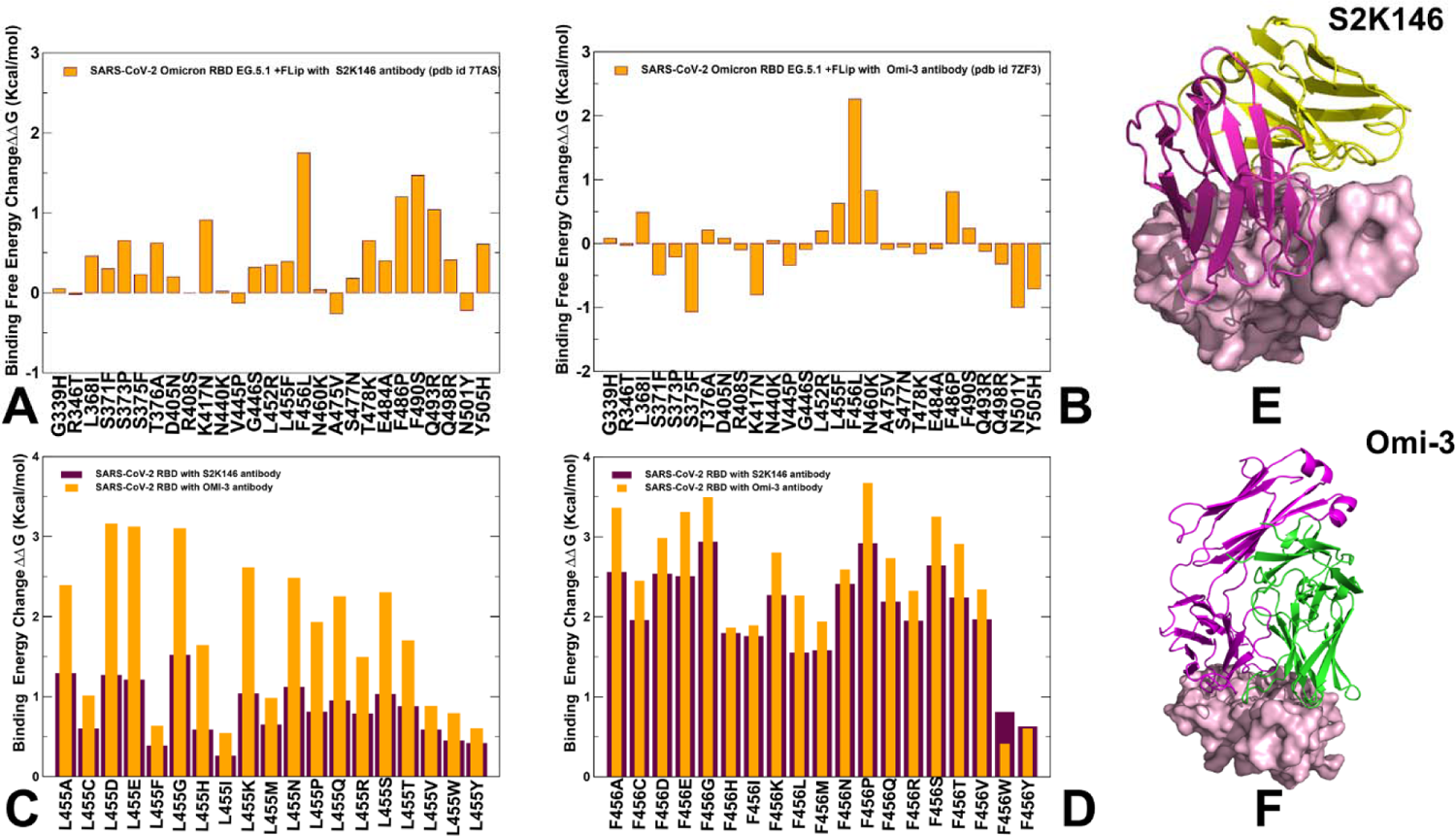
Structure-based mutational profiling of the S-RBD complexes with class 1 of RBD antibodies. The mutational profiling evaluates the binding energy changes induced by XBB.1.5 mutations in the RBD-antibody complexes. Mutational profiling of the S-RBD complex with S2K146 (A) and S-RBD Omicron complex with Omi-3 antibody (B). The binding free energy changes are shown in orange filled bars. The positive binding free energy values ΔΔG correspond to destabilizing changes and negative binding free energy changes are associated with stabilizing changes. (C) Mutational scanning of L455 residue in the RBD-S2K146 complex (maroon bars) and RBD-Omi-3 complex (orange bars). (D) Mutational scanning of F456 residue in the RBD-S2K146 complex (maroon bars) and RBD-Omi-3 complex (orange bars). The experimental structures of the RBD-antibody complexes are shown for RBD-S2K146 (E), RBD-Omi-3 (F). The RBD is shown in pink-colored surface representation and the antibodies are shown in ribbons (heavy chain in magenta and light chain in green-colored ribbons).

In the analysis of mutational scanning, we specifically focused on the S-antibody binding energy changes induced by XBB.1.5 mutations and L455F, F456L modifications (Figure 12). The binding free energy changes in the S-RBD complex with S2K146 (Figure 12A) showed appreciable losses of binding upon F456L, F486P and F490S mutations revealing that these substitutions are deleterious for S2K146 binding. At the same time, L455F induced only modest destabilization change (Figure 12A). For Omi-3 antibody, we found that F456 position presents the dominant binding affinity hotspot as the destabilization effect of F456L mutation (ΔΔG = 2.26 kcal/mol) is significantly stronger than for the S2K146 antibody (Figure 12B). Common to both S2K146 and Omi-3 antibodies, we also observed a considerable loss of binding due to F486P mutation (Figure 12A,B). Specific for Omi-3 antibody is a destabilizing role of N460K mutation inducing loss of binding ΔΔG =0.83 kcal/mol (Figure 12B). To examine further the effects of mutations in L455 and F456 positions on antibody binding, we also performed complete mutational scanning of these positions against S2K146 (Figure 12C) and Omi-3 (Figure 12D). We observed two interesting trends in this analysis. First, all modifications of L455 and F456 positions result in appreciable destabilization changes and for both positions the loss of binding is more significant for Omi-3 antibody. Importantly, we also found that binding losses for both antibodies are markedly larger upon mutations in F456 position (Figure 12D), Furthermore, while both L455 and F456 sites correspond to the binding hotspots for Omi-3 antibody, F456 is far more significant than L455 for binding with S2K146. These findings are in excellent agreement with neutralization profiling assays showing that XBB.1.5+F456L and EG.5.1 lineages are strongly resistant against Omi-3, Omi-18 and Omi-42 antibodies but cause less considerable loss of binding for S2K146 antibody.^42^ Mutational scanning analysis of the XBB-antibody binding and experimental evidence suggested that rapidly surging EG.5 and EG.5.1 variants may have evolved to improve the immune escape against class 1 RBD antibodies by using F456L and F486P mutations but this effect does not seem to be uniform and some monoclonal antibodies, such as S2K146, could still bind fairly strongly with XBB variants.^42^

Together, the results of mutational scanning and binding calculations of with ACE2 and antibodies clarified the role of FLip mutations in balancing fitness requirements for strong ACE2 binding and robust antibody escape. Indeed, our results suggested that F456L alone may not have a significant effect on ACE2 binding but could instead mediate strong epistatic couplings with L455F and Q453 to amplify the favorable contributions of these residues in ACE2 interactions. Concurrently, F456L together with F486P mutation are central for mediating antibody resistance which is consistent with functional experiments showing that the decreased neutralization of EG.5.1 relative to XBB.1.5 is primarily driven by XBB.1.5-F456L mutation.^42,43,52^ We argue that emergence of EG.5 and FLip variants may have been particularly driven by expanding the scope of mutations with strong epistatic effects that provide synergy between improved ACE2 binding and broad neutralization resistance. The findings of our study provide important quantitative rationale to the latest experimental evidence that ACE2 binding can be synergistically modulated and amplified via epistatic interactions of physically proximal binding hotspots, including Y501, R498, Q493, L455 and F456 residues.^46,57,58^ The results of forward and reversed mutational scanning also supported a central mediating role of Q493 in strengthening binding interactions with the ACE2 receptor. We suggest that through epistatic couplings revealed in the binding computations and network-based community analysis, the XBB lineages can leverage and amplify the favorable effect of the binding energy hotspots (L455, F45, Y489, Q493, R498, Y501) on ACE2 binding while the cumulative contribution of other RBD positions may be balanced against their prominent role in invoking the immune evasion. Our findings support a hypothesis according to which the impact on ACE2 binding affinity is mediated through a small group of universal binding hotspots while the effect of immune evasion could be more variant-dependent and modulated through recruitment of different mutational sites in the adaptable RBD regions.^80^ In this mechanism, the synergistic couplings between binding affinity hotspots may allow for accumulation of diverse and functionally beneficial (but binding neutral) mutations in other positions to balance tradeoffs between immune evasion and ACE2 binding. The revealed patterns are reminiscent of direct evolution studies showing that enhanced protein stability can promote broader evolvability and promote emergence of beneficial mutations while retaining the stability of the protein fold.^135–137^ Structure-based mutational scanning of the RBD binding interfaces with representative class RBD antibodies characterized the role of L455F and F456L mutations in eliciting broad resistance to neutralization, confirming that F456L and F486P may act as primary drivers of immune escape while individually incurring rather moderate changes in ACE2 binding. The observed inter-dependence between binding affinity hotspots and antibody resistance substitutions that is manifested by the epistatic couplings of Y501, R498, Q493, L455 and F456 residues can facilitate antibody escape and affect the direction of virus evolution, with potential implications for vaccine design.^59^

## Conclusions

Atomistic level structural predictions and microsecond atomistic MD simulations provided a thorough characterization of the AI-generated conformational ensembles and identified important differences in conformational landscape of the XBB variants. Through ensemble-based mutational scanning of the RBD residues and rigorous MM-GBSA calculations of binding affinities, we identified binding energy hotspots and characterized molecular basis underlying epistatic couplings between convergent mutational hotspots. Consistent with the DMS experiments, the results of this analysis quantified the increased stabilization effect of Q493 hotspot in the FLip variant and synchronization in favorable contributions of L455F and F456L mutations confirming a coordinated epistatic effect between Q493 and L455F/F456L doubled mutation in the FLip variant. The binding affinity computations provided a quantitative insight into the mechanism underlying stronger binding of the XBB.1.5 FLip variant relative to XBB.1 and XBB.1.5 showing that the differences in the binding affinity are driven by cumulative contributions of energetically coupled L455F, F456L and Q493 hotspots, while universally important for binding Y489, R498 and Y501 sites provide the bulk of favorable binding interactions in all complexes. The study supports a hypothesis that the impact of the increased ACE2 binding affinity on viral fitness is more universal and is mediated through cross-talk between convergent mutational hotspots, while the effect of immune evasion could be more variant-dependent. Mutational profiling is combined with network-based community analysis of epistatic couplings showing that the Q493, L455 and F456 sites mediate stable communities at the binding interface and serve as mediators of non-additive couplings. We found that pairs of Omicron substitutions with strong epistasis tend to be spatially proximal, and form localized stable modules allowing for compensatory energetic changes. Finally, our study examined the effects of antibody escape by conducting mutational profiling of RBD complexes with class 1 monoclonal antibodies revealing significant role of F456L mutation as a mediator of epistatic couplings and hotspot of immune resistance. Our data also pointed to the key role of R493Q mutation in modulating affinity-enhancing effects of mutational changes in Y453, L455 and F456, showing that several key RBD hotspots (N501Y, Q498R and R493Q) can exploit variant-specific epistatic couplings with immune-escaping positions to protect ACE2 binding affinity with “minimum resources” while enabling mutational diversity to ensure broad immune evasion. This interpretation is consistent with the notion that functionally balanced substitutions that simultaneously optimize immune evasion and high ACE2 affinity may continue to emerge through lineages with beneficial pair or triplet combinations of RBD mutations involving mediators of epistatic couplings and sites in highly adaptable RBD regions.

## Data Availability Statement

Data is fully contained within the article and Supplementary Materials. All simulations were performed using the all-atom additive CHARMM36M protein force field that can be obtained from http://mackerell.umaryland.edu/charmm_ff.shtml. The rendering of protein structures was done with ChimeraX package (https://www.rbvi.ucsf.edu/chimerax/) and Pymol (https://pymol.org/2/). The software tools used in this study are freely available at GitHub sites: https://github.com/deepmind/alphafold; https://github.com/sokrypton/ColabFold/; https://github.com/RSvan/SPEACH_AF; https://github.com/Amber-MD/cpptraj. All the data obtained in this work (including simulation trajectories, topology and parameter files, the software tools, and the in-house scripts are freely available at ZENODO website https://doi.org/10.5281/zenodo.10310692; https://zenodo.org/records/10358578.

## Supporting Information

The post-processing AF2 analysis of predictions for the XBB.1 RBD residues in the RBD-ACE2 complex includes the pLDDT per RBD residue for the top five models and the predicted alignment error (PAE) for the top five models obtained from AF2 predictions. (Figure S1). Figure S2 presents the AF2 analysis of predictions for the XBB.1.5 RBD residues in the RBD-ACE2 complex and includes the pLDDT per RBD residue for the top five models and the predicted alignment error (PAE) for the top five models obtained from AF2 predictions. Figure S3 presents the distributions of the GDT_TS structural similarity metric for the XBB RBD-ACE2 conformational ensembles obtained from AF2-MSA depth predictions. Figure S4 presents the distributions of the RMSD structural similarity metric for the XBB RBD-ACE2 conformational ensembles obtained from AF2-MSA depth predictions. Figure S5 shows the scatter plots of the DMS-based binding free energy changes for the RBD residues and computational mutational scanning of the RBD residues for the BA.1, BA.2 and XBB.1.5 RBD-ACE2 complexes. Figure S6 presents the residue-based decomposition of the MM-GBSA energies for the XBB.1.5 +L455F RBD-ACE2 complex and XBB.1.5+F456L RBD-ACE2 complex. Figure S7 describes the results of network-based community decomposition for the binding interface residues in the XBB RBD-ACE2 complexes. This material is available free of charge via the Internet at http://pubs.acs.org.

## Author Contributions

Conceptualization, G.V.; methodology, N.R., M.A., G.C., S.X., G.V. P.T.; software, N.R., S.X., M.A., G.G., G.V and P.T.; validation, N.R., G.V.; formal analysis, N.R., G.V., M.A., G.G., S.X., and P.T.; investigation, N.R., M.A., G.C., G.V. and P.T.; resources, N.R., G.V., M.A. S.X., and G.V.; data curation, N.R., M.A., G.C.,G.V.; writing— original draft preparation, N.R., M.A., G.V.; writing—review and editing, N.R. and G.V.; visualization, N.R., M.A., G.C., S.X., G.V. G.V.; supervision, G.V.; project administration, G.V.; funding acquisition, P.T. and G.V. All authors have read and agreed to the published version of the manuscript.

