## Supplemental Figures S1-S7 for "AlphaFold2-Enabled Atomistic Modeling of Epistatic Binding Mechanisms for the SARS-CoV-2 Spike Omicron XBB.1.5, EG.5 and FLip Variants: Convergent Evolution Hotspots Cooperate to Control Stability and Conformational Adaptability in Balancing ACE2 Binding and Antibody Resistance"

### Supplementary Materials

#### AI-Augmented Atomistic Modeling of Structural Ensembles and Receptor Binding for the SARS-CoV-2 Spike Omicron XBB Lineages: Convergent Evolution Hotspots in the XBB.1.5, EG.5 and FLip Variants Exploit Epistatic Couplings for Optimizing Balance of ACE2 Binding and Antibody Escape

Nishank Raisinghani, [1] Mohammed Alshahrani,[1] Grace Gupta,[1] Sian Xiao[3], Peng Tao[3], Gennady Verkhivker[1,2]*

Keck Center for Science and Engineering, Graduate Program in Computational and Data Sciences, Schmid College of Science and Technology, Chapman University, Orange, CA 92866, United States of America

[2] Department of Biomedical and Pharmaceutical Sciences, Chapman University School of Pharmacy, Irvine, CA 92618, United States of America

[3]Department of Chemistry, Center for Research Computing, Center for Drug Discovery, Design, and Delivery (CD4), Southern Methodist University, Dallas, Texas, 75275, United States of America

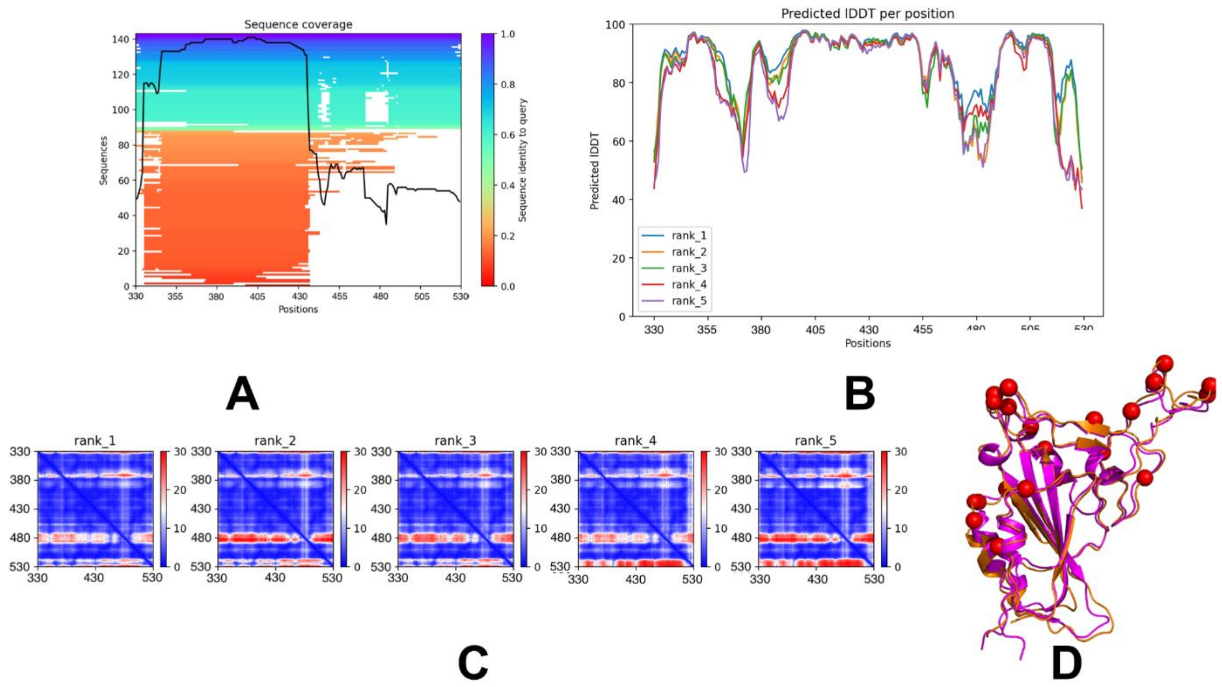


**Figure S1.** The post-processing AF2 analysis of predictions for the XBB.1 RBD-ACE2 complex. (A) A heatmap representation of the MSA indicates all sequences mapped to the input sequences. The color scale points to the identity score, and sequences are ordered from top (largest identity) to bottom (lowest identity). White regions are not covered, which occurs with sub-sequence entries in the database. The black line qualifies the relative coverage of the sequence with respect to the total number of aligned sequences. (B) The pLDDT per RBD residue for the top five models obtained from AF2 predictions of the XBB.1 RBD-ACE2 complex. (C) The predicted alignment error (PAE) for the top five models obtained from AF2 predictions. These heat maps are provided for each final model and show the PAE between each residue in the model. The color scale contains three colors to highlight the contrast between the high confidence regions and the low confidence regions.

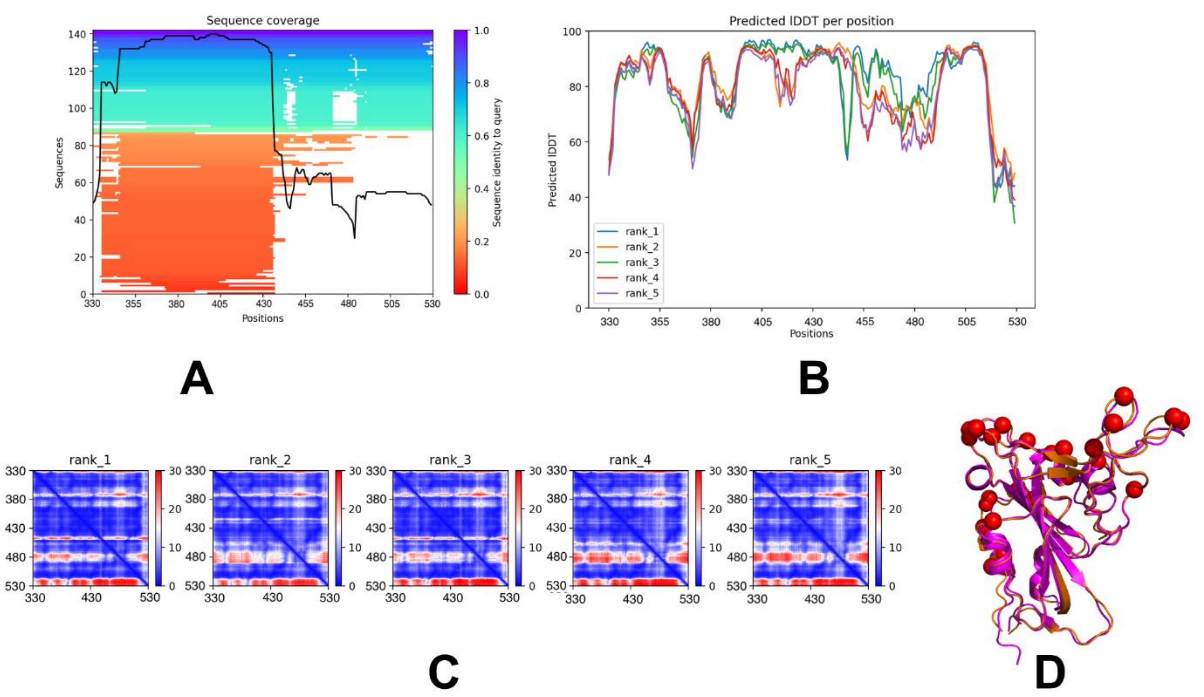


**Figure S2.** The post-processing AF2 analysis of predictions for the XBB.1.5 RBD-ACE2 complex. (A) A heatmap representation of the MSA indicates all sequences mapped to the input sequences. The color scale points to the identity score, and sequences are ordered from top (largest identity) to bottom (lowest identity). White regions are not covered, which occurs with sub-sequence entries in the database. The black line qualifies the relative coverage of the sequence with respect to the total number of aligned sequences. (B) The pLDDT per RBD residue for the top five models obtained from AF2 predictions of the XBB.1.5 RBD-ACE2 complex. (C) The predicted alignment error (PAE) for the top five models obtained from AF2 predictions. These heat maps are provided for each final model and show the PAE between each residue in the model. The color scale contains three colors to highlight the contrast between the high confidence regions and the low confidence regions.

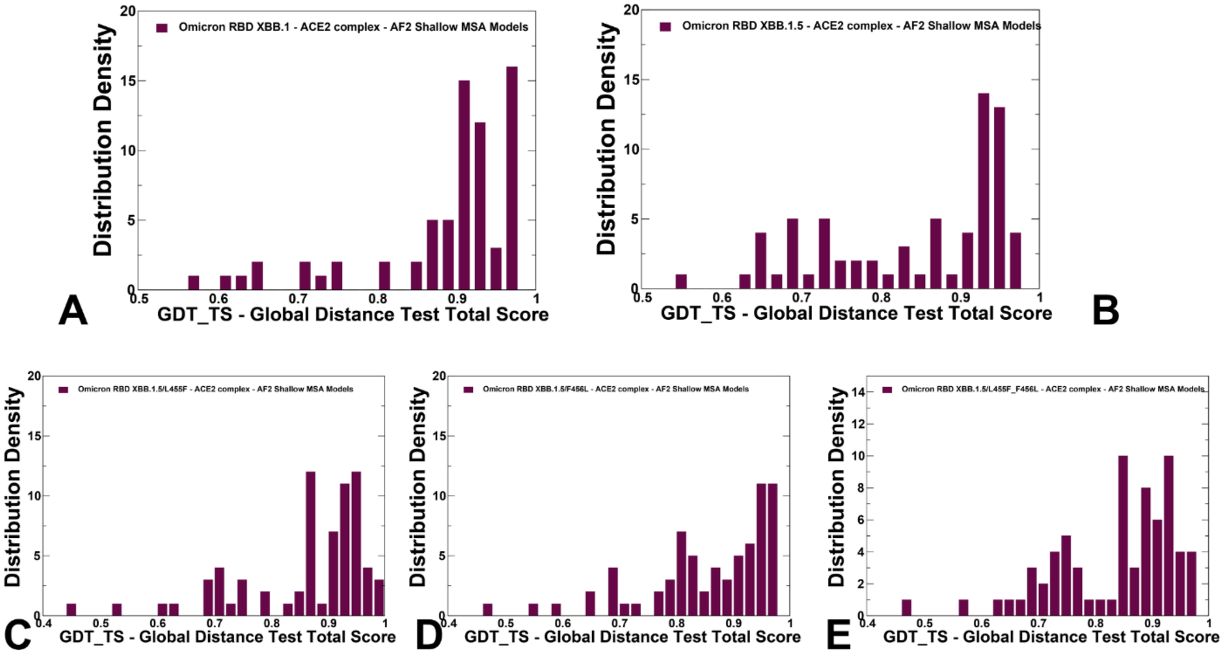


**Figure S3.** The density distribution of the GDT_TS score measuring structural similarity of the predicted RBD conformational ensembles with respect to the structure of the XBB.1.5 RBD-ACE2 (pdb id 8WRL) for XBB.1 (A) XBB.1.5 (B), XBB.1.5+L455F(C), XBB.1.5+F456L (D) and XBB.1.5+L455F/F456L FLip variants (E).

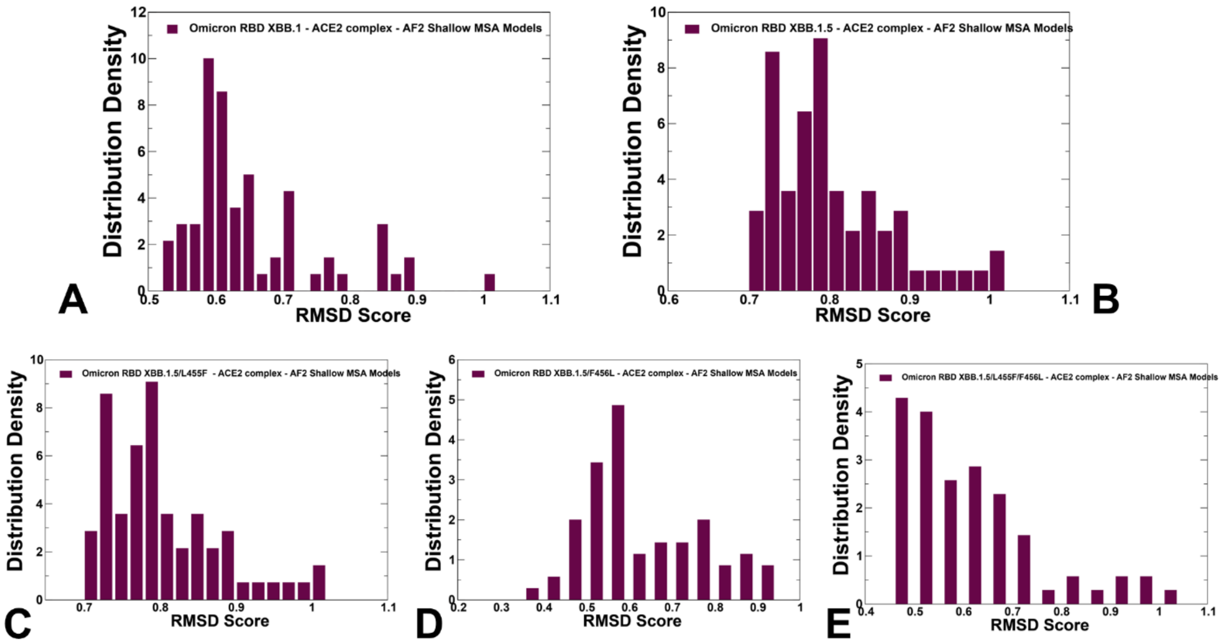


**Figure S4.** The density distribution of the RMSD score measuring structural similarity of the predicted RBD conformational ensembles with respect to the structure of the XBB.1.5 RBD-ACE2 (pdb id 8WRL) for XBB.1 (A) XBB.1.5 (B), XBB.1.5+L455F(C), XBB.1.5+F456L (D) and XBB.1.5+L455F/F456L FLip variants (E).

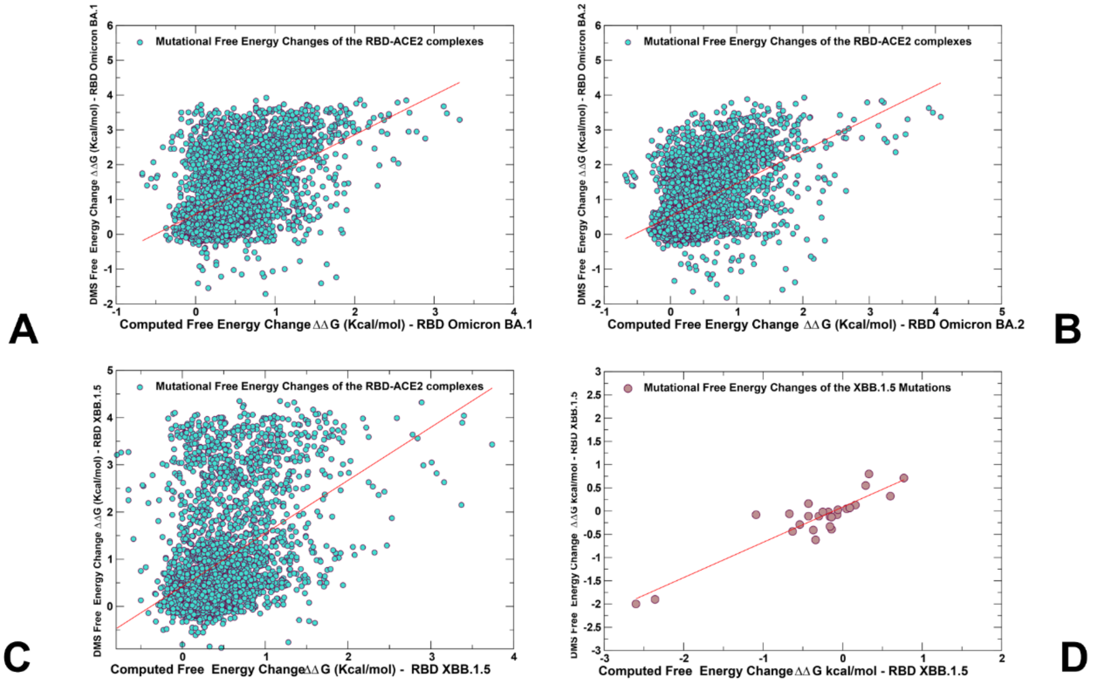


**Figure S5.** The scatter plots of the DMS-derived binding free energy changes for the RBD residues and computational mutational scanning of the RBD residues to estimate mutational effects on ACE2 binding. The effect on ACE2 receptor-binding affinity ($\Delta$ log10 $K_D$) of every single amino-acid mutation in SARS-CoV-2 RBD was experimentally determined by high-throughput titration assays using DMS experiments.[54-56,62] The results of computational mutational scanning of the RBD residues were averaged over conformational ensembles obtained from all-atom MD simulations. The scatter plot of the experimental and computed binding free energy changes from mutational scanning of the RBD residues in the Omicron BA.1 RBD-ACE2 complex, pdb id 7WBP (A) , BA.2 RBD-ACE2 complex, pdb id 7XB0 (B) and XBB.1.5 RBD-ACE2 complex, pdb id 8WRL (C). The correlation coefficient R=0.65 for BA.1 RBD, R=0.68 for BA.2 RBD and R=0.52 for XBB.1.5 RBD (D) The scatter correlation between experimental and computed binding free energies for XBB.1.5 mutational sites only. The data points are shown in light-brown colored circles.

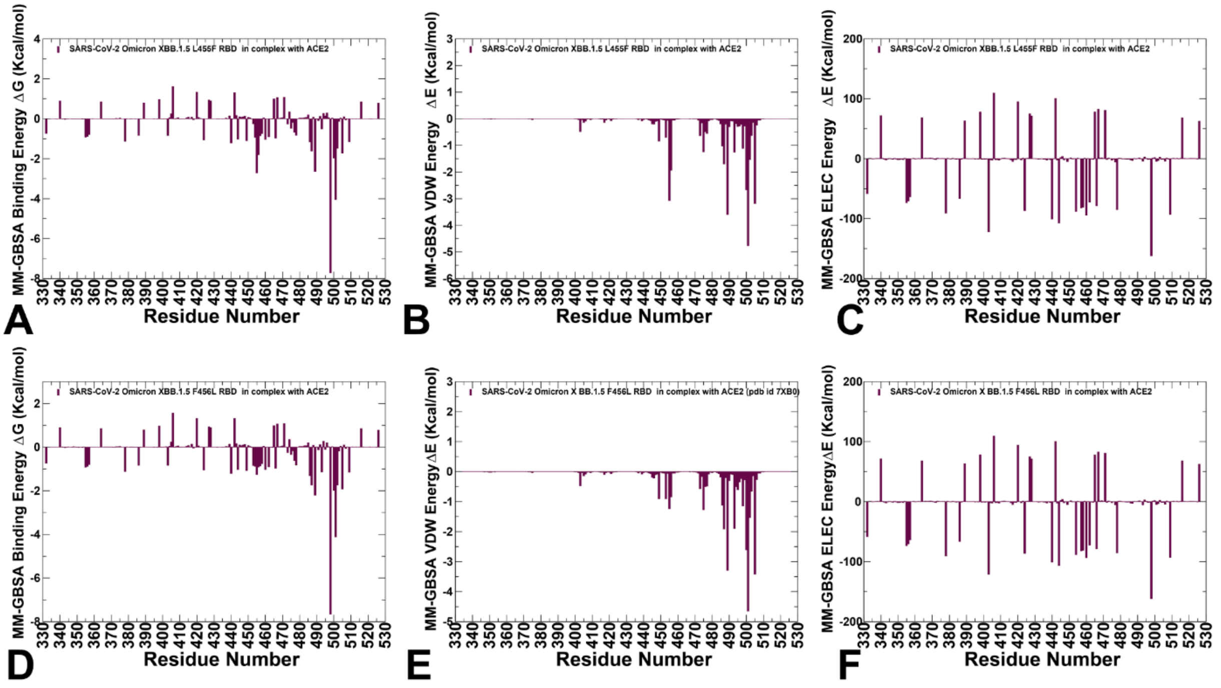


**Figure S6.** The residue-based decomposition of the MM-GBSA energies. (A) The residue-based decomposition of the total MM-GBSA binding energy ΔG contribution for the XBB.1.5+L455F RBD-ACE2 complex. (B) The van der Waals contribution of the total binding energy for the XBB.1.5+L455F RBD-ACE2 complex. (C) The electrostatic contribution of the total binding energy for the XBB.1.5+L455F RBD-ACE2 complex. The residue-based decomposition of the total MM-GBSA binding energy ΔG contribution for the XBB.1.5+F456L RBD-ACE2 complex (D) , the van der Waals contribution of the XBB1.5+F456L total binding energy (E) and the electrostatic contribution of the XBB.1.5+F456L total binding energy (F). Th MM-GBSA contributions are evaluated using 1,000 samples from the equilibrium MD simulations of respective RBD-ACE2 complexes. It is assumed that the entropy contributions for binding are similar and are not considered in the analysis. The statistical errors was estimated on the basis of the deviation between block average and are within 0.25-0.95 kcal/mol.

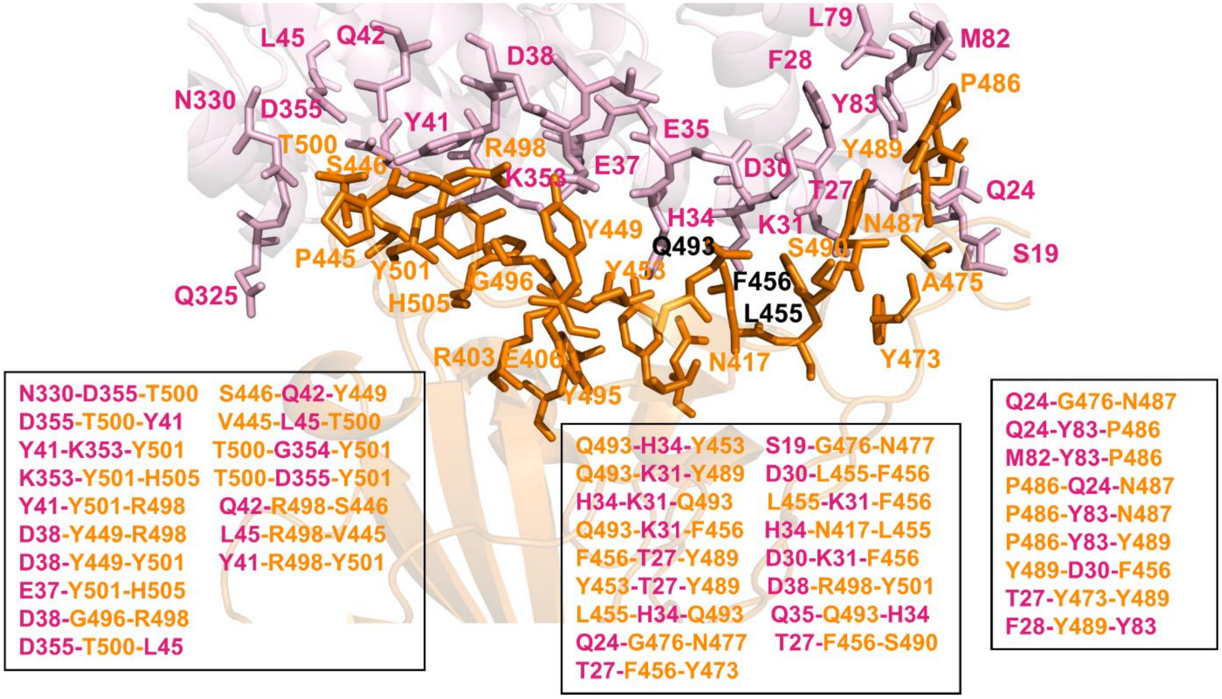


**Figure S7.** Structural mapping and full annotation of the intermolecular 3-cliques for the XBB.1.5 RBD-ACE2 complex. The RBD binding interface residues are shown in orange sticks and ACE2 binding interface residues are shown in pink sticks.
